# Independence and Coherence in Temporal Sequence Computation across the Fronto-Parietal Network

**DOI:** 10.1101/2025.08.25.672149

**Authors:** Hiroto Imamura, Fumiya Imamura, Reiko Hira, Yoshikazu Isomura, Riichiro Hira

**Author notes:** Corresponding authors. Correspondence: Riichiro Hira, M.D. Ph.D. Department of Physiology and Cell Biology, Graduate School of Medical and Dental Sciences, Institute of Science Tokyo, 1-5-45 Yushima, Bunkyo-ku, Tokyo, 113-8510, Japan. Yoshikazu Isomura, Ph.D. Department of Physiology and Cell Biology, Graduate School of Medical and Dental Sciences, Institute of Science Tokyo, 1-5-45 Yushima, Bunkyo-ku, Tokyo, 113-8510, Japan.

## Abstract

Time processing requires distributed and coordinated cortical dynamics. Flexible yet robust temporal representations can arise from two distinct computational modes: a coherence mode, where multiple cortical areas hold the same elapsed-time estimate, and an independence mode, where each area maintains its own local estimate. However, how the brain switches between these modes has remained unknown. Using mesoscale two-photon calcium imaging, we simultaneously recorded neuronal populations in the secondary motor cortex (M2) and posterior parietal cortex (PPC) of mice performing a novel alternating-interval timing task. Both areas encoded elapsed time through similar high-dimensional sequential activity. Decoding analyses revealed that the fronto-parietal network has both independent and coherent temporal codes. Communication-subspace analysis showed that temporal information was distributed across multiple low-variance subspaces, whereas the largest subspace preferentially encoded behaviour. A twin recurrent neural network (RNN) model with sparse inter-RNN connections and shared high-variance noise reproduced these experimental findings. Moreover, perturbations applied along the dominant shared subspace paradoxically enhanced independence between the two networks. Through a mathematical formalization based on the local Lyapunov exponents, we uncovered how perturbations along different subspaces selectively evoke either independent or coherent communication mode. Together, these results reveal a principle by which fronto-parietal circuits achieve robust yet flexible computation through the interplay of sparse coupling and shared global fluctuations.

## Introduction

Temporal information processing underlies a wide spectrum of cognitive functions (Paton and Buonomano 2018; Buonomano and Laje 2010; Tsao et al. 2022). Neural representations of time have been widely observed in the prefrontal cortex, **secondary motor cortex (M2)**, **posterior parietal cortex (PPC)**, and in subcortical structures such as the basal ganglia and cerebellum (Murakami et al. 2014; Shimbo et al. 2024; Buhusi and Meck 2005; Petter, Gershman, and Meck 2018; Pastalkova et al. 2008; MacDonald et al. 2011; Kraus et al. 2013; Gouvêa et al. 2015; Mello, Soares, and Paton 2015; Bakhurin et al. 2017; Toso et al. 2021; Bolkan et al. 2017; Tiganj et al. 2017; Mita et al. 2009; Kalmbach et al. 2010; Simen et al. 2011; Johansson et al. 2014; Hira et al. 2024; Jin, Fujii, and Graybiel 2009). Elapsed time has traditionally been observed as ramp-up or ramp-down patterns of neural activity, while sequential activity, where neurons that peak at intermediate times fire in succession, is also widespread (Harvey, Coen, and Tank 2012; Hira et al. 2024; Mita et al. 2009). Modelling studies have reproduced such sequences using recurrent neural networks (RNNs) (Rajan, Harvey, and Tank 2016; Pereira and Brunel 2019; Zhou, Masmanidis, and Buonomano 2020; Wilson et al. 2024; Wang et al. 2017; Hardy et al. 2018).

While RNNs can reproduce temporal processing within a single brain area, it remains unknown how multiple areas coordinate or decouple to flexibly switch between these distinct coding modes. Because time-representing areas are thought to be linked functionally, their temporal representations might be globally synchronized (Paton and Buonomano 2018; Shimbo et al. 2024; Tang, Shin, and Jadhav 2021). Conversely, when each area must track the elapsed time from different events, their activity could become asynchronous. When multiple brain areas share temporal information, it is thought that they must flexibly switch between a **coherence mode**, in which all regions operate on a shared clock, and an **independence mode**, in which each region maintains its own local clock. Crucially, most existing models omit key biological constraints such as shared low-frequency noise and sparse long-range excitation, limiting their correspondence with experimental observations. A biologically grounded multi-network framework is therefore required to close this gap between theory and experiment (Huang et al. 2019; Kleinman, Chandrasekaran, and Kao 2021; Mastrogiuseppe and Ostojic 2018; Voytek et al. 2015; Seeman et al. 2018; Clark and Beiran 2025).

M2 and PPC are reciprocally connected and are known to support various cognitive functions including working memory and sensory prediction (Voitov and Mrsic-Flogel 2022; Raltschev et al. 2025; Olsen et al. 2019; Hovde et al. 2019; Hwang, Sato, and Sato 2021; Barthas and Kwan 2017). In working-memory tasks, recurrent cortico-cortical connections between M2 and PPC are essential for maintaining the memory trace (Voitov and Mrsic-Flogel 2022). Interestingly, the dominant component of neural activity shared by M2 and PPC was unrelated to working memory (Voitov and Mrsic-Flogel 2022). The principal dynamics shared by a pair of cortical areas may also be shared across many regions and have been suggested to relate to motor information (MacDowell et al. 2025; Hira et al. 2024; Stringer et al. 2019). Anatomically, M2 and PPC exhibit a distinctive organization characterized by sparse long-range excitatory projections and shared inputs from widespread sources (Olsen et al. 2019; Hovde et al. 2019; Zhang et al. 2016; Hooks et al. 2013; Oh et al. 2014), features that are theoretically well suited for modulating the switch between independence and coherence. Moreover, the two areas are on the dorsal cortical surface of the mouse brain, making them ideally positioned for simultaneous large-field two-photon imaging and direct comparison between experimental data and computational models.

Here we combine behaviour, mesoscale imaging, and modelling to uncover how the M2–PPC network balances coherent and independent temporal coding. Specifically, we (1) developed a novel alternating-interval task in which 6-s and 12-s intervals appear alternately, (2) simultaneously recorded thousands of neurons in M2 and PPC using a large field-of-view two- photon microscope (Hira et al. 2024; Yu et al. 2021), and (3) constructed and analysed a twin RNN model using the acquired data to empirically examine the dynamic balance shaped by sparse connectivity and global noise. Our results support a two-factor mechanism in which sparse inter-areal excitation promotes coherence, whereas shared 1/f-type global noise Favors independence. More broadly, the seemingly redundant coding distributed across multiple brain areas may reflect a trade-off between robustness for ensuring stable computation and flexibility for supporting a rich computational repertoire that adapts to diverse demands in natural environments.

## Result

### Development of an alternating-interval task

M2 and PPC dynamically maintain a representation of continuous time and update their internal states in response to external inputs (**Fig. 1a**). This capability persists even when successive trials are separated by a long inter-trial interval (Akrami et al. 2018). To probe M2 and PPC effectively, we need a temporally continuous task with no distinction between individual trials, such that the animal must continuously generate internal states based on the current state and external input. From this perspective, we compared several candidate paradigms, including previously used designs (Zhou, Masmanidis, and Buonomano 2020; Toda et al. 2017; Mello, Soares, and Paton 2015), to identify the simplest task satisfying these requirements (**Fig. 1b**). **Task-1** delivers a cue and reward at a fixed 6-s interval. Although the animals would spontaneously learn to anticipate the 6-s delay, the new reward alone could inform the task state without referring to the internal state. Thus, continuous updating of internal states is not necessary. **Task-2** presents two cues, cue1 and cue2, that predict reward 6 s and 12 s later, respectively. Again, each trial is independent, resulting in representational discontinuities across trials when the cue is presented. **Task-3** is a modified delayed match/non-match paradigm involving two cue types. In each trial, the cue and reward were presented simultaneously, either 6 or 12 seconds apart. A 6-second interval was used when the current cue matched the previous one; otherwise, a 12-second interval was applied. Because the predicted interval depends on the conjunction of working memory and the current cue, internal representations remain continuous through trials. Thus, this task is well suited for investigating M2 and PPC, although its complexity may require an extensive training period.

**Figure 1.**
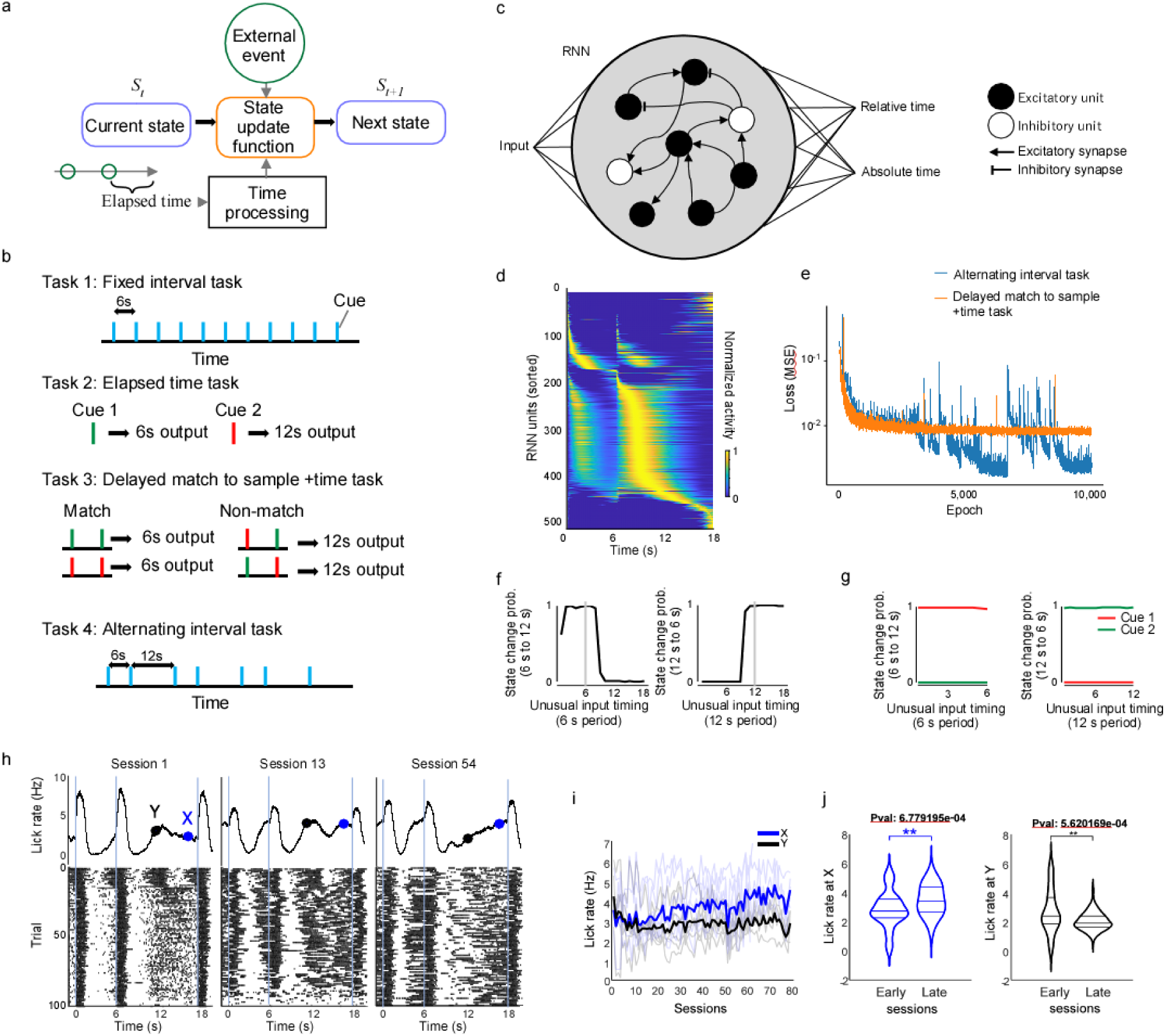
Alternating-Interval (AI) task for investigation of fronto-parietal network function **a.** Hypothetical computational roles of M2 and PPC explored in this study. **b.** Four candidate tasks. For all tasks, the subjects need to predict the timing of the next reward. In the fixed-interval task (Task 1), a cue or reward is delivered every 6 s. In the elapsed-time task (Task 2), Cue 1 and Cue 2 signal rewards 6 s and 12 s later, respectively. In the DMS-time task (Task 3), if the current cue matches the previous cue, a reward follows 6 s later, otherwise the reward follows after 12 s. In the alternating-interval task (Task 4), the intervals between cues or rewards alternate between 6 s and 12 s. **c.** Schematic of the RNN. The RNN consists of 80 % excitatory and 20 % inhibitory units and is trained to output absolute and relative time until the next reward. **d.** Trial-averaged RNN activity after training on the AI task, sorted and normalized by the time of maximum activity. **e.** Examples of loss during training on the alternating-interval task and the delayed match-to-sample + time task. **f.** Immediately after an unusual input timing, activities of the RNN units were decoded to determine whether the activity pattern was closer to the 6-s-interval trajectory or the 12-s-interval trajectory. For unusual inputs delivered during a 6-s interval, the probability of the decoding output switching to the 12-s trajectory within the following 1 s was plotted as a function of the input interval(left). Similarly, for unusual inputs delivered during a 12-s interval, the probability of the decoding output switching to the 6-s trajectory within the next 1 s was plotted against the input time (right). **g.** In the elapsed-time task, the probabilities of decoding output switching after unusual inputs were plotted in the same manner as in **f**. Blue corresponds to Cue 1 (6-s output) and red to Cue 2 (12-s output). **h.** Behavioural changes in a representative mouse (mouse #2) during the AI task. In session 3, even during 12-s trials the lick rate rose until 6 s and then plateaued. In session 17, during 12-s trials the lick rate decreased after 6 s before increasing again. By session 64, the lick rate in 12-s trials increased continuously. **i.** Changes in lick rates 5 s (Y, black) and 11 s (X, blue) into 12-s trials from the start of AI task training. **j.** Differences in lick rates X and Y between early sessions (1–10) and late sessions (20–80) (*p < 0.05, **p < 0.01, Wilcoxon rank-sum test).

We therefore designed an **alternating-interval task (AI task, task-4)**. In this paradigm, a cue and reward are delivered at fixed intervals that alternate between 6 s and 12 s. Consequently, when a cue appears, the animal must maintain its internal state depending on the duration of the preceding wait in order to predict the next interval. Because cue discrimination is unnecessary, this task should be easier for mice to learn than task-3. For clarity, we define the **6-s interval** and the **12-s interval** within a continuous 18-s epoch: the 6-s interval starts at trial time 0 s, the 12-s interval begins at 6 s, and the epoch ends at 18 s.

To anticipate how the brain might respond to tasks-2, -3, and -4, we trained recurrent neural networks (RNNs) to output both absolute and relative elapsed time of the target durations (**Fig. 1c; Extended Data** Fig. 1). In task-4, the RNN agent achieved correct performance and, after learning, generated sequential activity (**Fig. 1d**). The agent also succeeded on task-2, but our simple RNN failed to learn task-3 (**Fig. 1e**) We next compared tasks-2 and -4 by probing the networks with unusual inputs. In task-2, presentation of cue1 or cue 2 always evoked the 6-s or 12-s trajectory, respectively, regardless of when the unusual input occurred (**Fig. 1f**). In task-4, however, a cue delivered before 9 s in the 6-s period drove a transition to the 12-s trajectory, and a cue delivered after 9 s within the 12-s period drove a transition to the 6-s trajectory (**Fig. 1g**). Thus, unlike task-2, task-4 promotes a state-dependent mechanism in which internal dynamics are updated by both elapsed time and external events. These features make the task-4, AI task, well suited for dissecting M2 and PPC functions. Although the paradigm could be implemented as an operant task, we used a classical-conditioning version in this study.

### Mice learn AI task

Can head-fixed mice learn the AI task? We trained water-restricted mice with head-fixed conditions. During pre-training, the reward interval was fixed at 6 s. Subsequently, the reward intervals were alternated between 6 and 12 s to encourage the mice to predict the upcoming interval from the history of previous intervals. As training progressed in the pre-training (fixed-interval) task, the mice showed predictive licking peaking at 6 s after each reward. Once this anticipatory licking emerged, the task was switched to the AI task. Early in AI-task training, the mice still showed anticipatory licking that peaked at 6 s even during 12-s intervals (**Fig. 1h**; session 3). As training progressed, they began to display anticipatory licking that peaked at 12 s during 12-s intervals (**Fig. 1h**; sessions 17 and 64). At the same time, during 6-s intervals they continued to lick predictively with a peak at 6 s. The lick rate at 5 s into the 12-s interval quickly declined and reached a plateau after five sessions, whereas the lick rate at 11 s gradually increased as training advanced (**Fig. 1i, j**; n = *5* mice). Thus, in the AI task, the mice learned that 6-s and 12-s intervals alternate.

### Single neuron activity was similar in M2 and PPC

We examined population activity in M2 and PPC using two-photon calcium imaging with a Diesel2p mesoscope after the mice mastered the task (**Fig. 2a, b**; n = 5 mice). Both M2 and PPC can be captured simultaneously within the FOV of the Diesel2p. We broadly expressed GCaMP in layer 2/3 of M2 and PPC either by local AAV injections (n = 5 mice, **Extended Data** Fig. 2) or by neonatal intracerebral injections performed on postnatal day 1 (**Fig. 2a, b**). We functionally identified sensory areas using one-photon calcium imaging and defined M2 and PPC accordingly (**Extended Data** Fig. 2). We obtained 1,463 ± 418 neurons in M2 and 815 ± 590 neurons in PPC (6 sessions from 5 mice).

**Figure 2.**
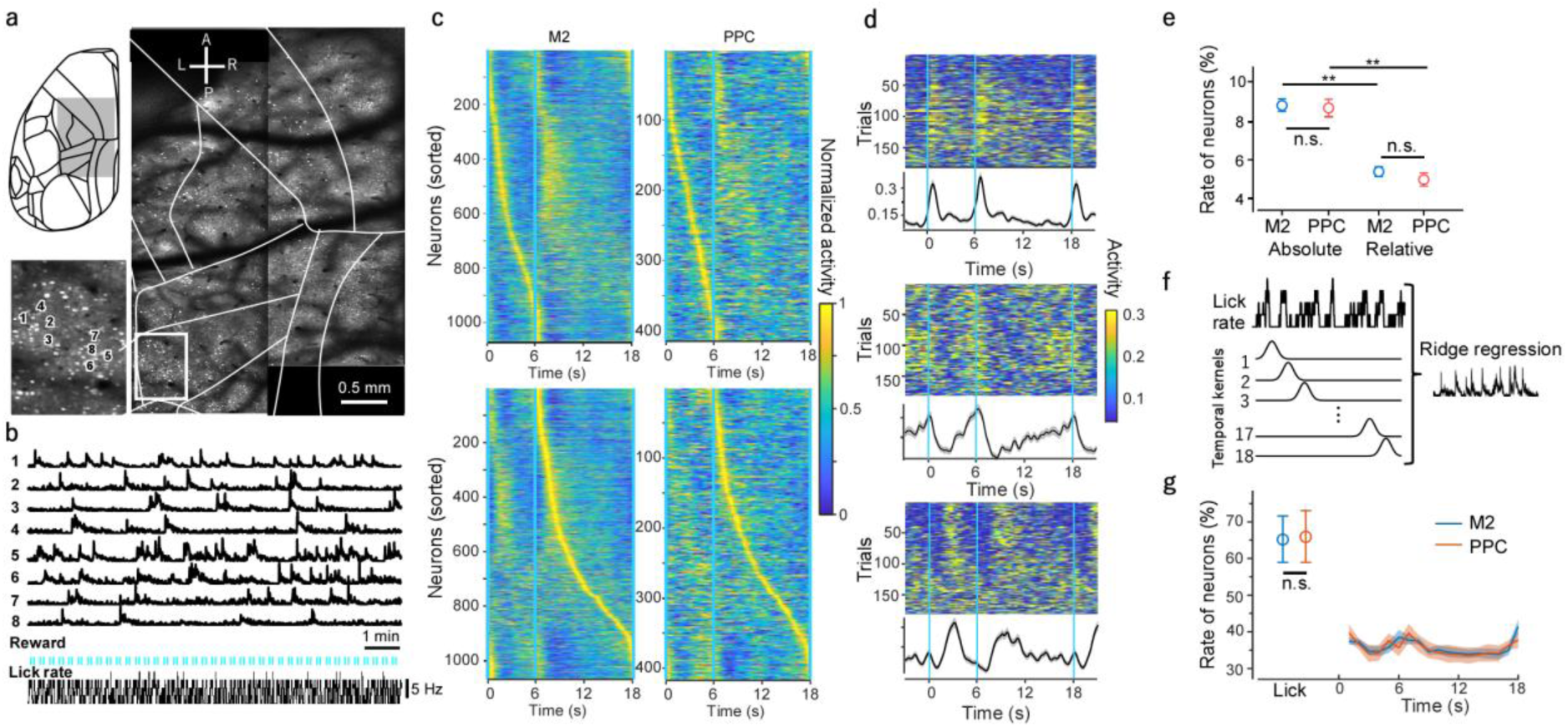
Large field-of-view two photon calcium imaging during the AI task. a. Imaging field capturing M2 and PPC simultaneously. **b.** Eight representative neuronal traces alongside licking behaviour and reward timing. **c.** Trial-averaged activity of simultaneously recorded M2 and PPC neurons; neurons are sorted by peak time in the 6-s interval (upper) and the 12-s interval (lower). **d.** Three representative single-neuron activity patterns. **e.** Proportions of neurons whose peak activity reflect absolute time (left) versus relative time (right) (**p <0.01, Wilcoxon rank-sum test). **f.** Diagram of the encoding model (ridge regression). **g.** Proportion of neurons whose activity was significantly explained by the lick-rate or time-window predictors; no significant difference between M2 and PPC (lick rate: p =0.84, Wilcoxon rank-sum; time window: p = 0.90, ANOVA).

For each neuron in M2 and PPC, we visualized the mean activity during the 6-s and 12-s intervals sorted by peak time (**Fig. 2c**). In both intervals the peak distribution covered the entire duration, exhibiting sequential activity. Among them were neurons active immediately after reward (**Fig. 2d, top**), neurons whose activity scaled between the 6-s and 12-s intervals (**Fig. 2c, middle**; temporal-scaling neurons), and neurons that peaked several seconds after reward (**Fig. 2c, bottom**). Neurons encoding absolute time accounted for 9.25 % in M2 and 8.85 % in PPC, whereas temporal-scaling neurons encoded relative time at 5.49 % in M2 and 5.91 % in PPC. There was no significant difference between M2 and PPC (M2: p = 0.81, PPC: p = 0.35, permutation test), but neurons that encoded absolute time were more prevalent in both regions (M2: p < 1 × 10⁻³, PPC: p < 1 × 10⁻³, permutation test; **Fig. 2e**). To account for the diverse neuronal activities, we constructed an encoding model using a linear model with L2 norm regularization (Ridge regression, **Fig. 2f)**. The explanatory variables were lick rate and elapsed time within each trial. A systematic analysis revealed comparable proportions of neurons related to lick rate and elapsed time in M2 and PPC (**Fig. 2g**). These results indicate that individual neurons in M2 and PPC exhibit similar task relevance and possess comparable timing representations.

### Similar sequential dynamics exhibited by population activity in M2 and PPC

Next, we investigated temporal representation at the population level. First, to determine how the populations in M2 and PPC displayed sequential dynamics, we compared peak entropy, temporal sparsity, and the sequentiality index between them (Methods, **Extended Data** Fig. 3a). No significant differences were found between M2 and PPC for any of these metrics. These results suggest that the dynamics in both areas are equally sequential. In populations with sequential dynamics, neurons that share similar temporal preferences may receive common inputs, increasing noise correlations. In M2, we found that neuron pairs whose peaks were close during the 6-s interval had stronger correlations of trial-to-trial fluctuations during the 12-s interval (**Extended Data** Fig. 3b). Likewise, pairs with nearby peaks during the 12-s interval showed larger correlations of fluctuations during the 6-s interval. The same relationships were observed in PPC (**Extended Data** Fig. 3c). Moreover, they were also significant for pairs containing one M2 neuron and one PPC neuron (**Extended Data** Fig. 3d). These findings indicate that neurons with similar temporal preferences in M2 and PPC interact, either synaptically or via common inputs.

### Both M2 and PPC exhibit high-dimensional representations of time

To investigate how neuronal populations in M2 and PPC maintain temporal information, we first visualized their activity with UMAP (**Fig. 3a**). In both areas the trajectories formed an 18- s circular loop. We then compared these temporal representations by decoding elapsed time. Because the number of recorded neurons was large, we first restricted the analysis to subspaces rich in temporal information using contrastive PCA (cPCA, see **Supplementary information**, **Fig. 3b; Extended Data** Fig. 4). In cPCA, we prioritized dimensions with high within-trial variance corresponding to the 18-s trial activity, and low between-trial variance reflecting consistency across trials. The trade-off between these variances was governed by the parameter 𝑤, which we chose to maximize average decoding accuracy across animals (𝑤 = 0.60 for M2 and 0.50 for PPC; **Extended Data** Fig. 4a, b). We decoded time with a random-forest classifier. Both regions achieved moderate accuracy (**Fig. 3c**) without a significant difference between them (**Fig. 3d**). Decoding accuracy depended on the degree of dimensionality reduction (**Fig. 3e**), yet the dimensionality yielding peak performance was similar in the two regions (∼15–50; **Fig. 3f**). These results show that M2 and PPC encode time in comparably high-dimensional subspaces, relying on broadly distributed neuronal activity with little difference in representational format.

**Figure 3.**
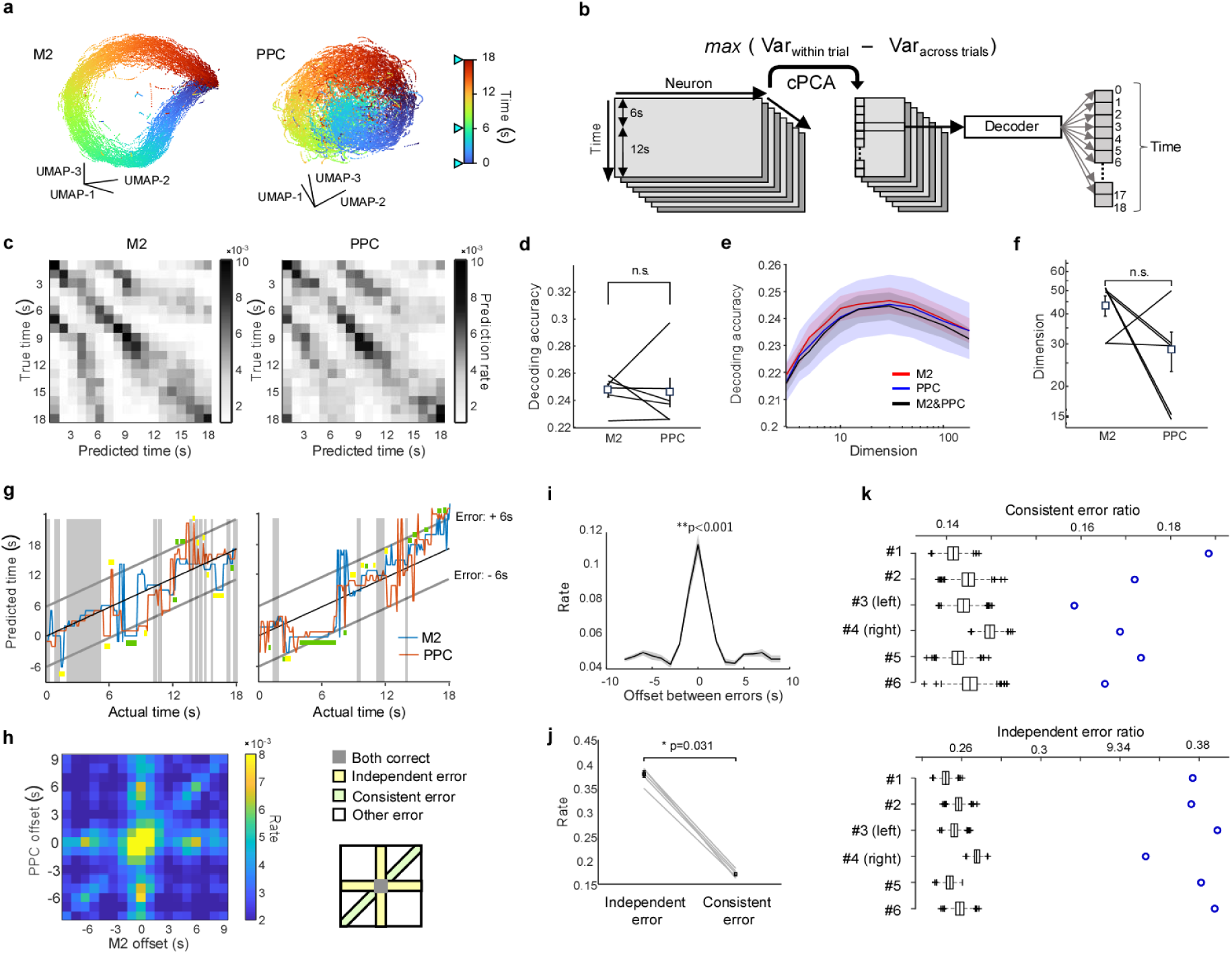
Decoding analysis on the neural data. a. Example neural manifold analysis. Population activity of a single session was visualized with cPCA (w=0.6), followed by 3d projection using UMAP. **b.** Schematic of time decoding analysis. The dimension of the population activity was reduced with cPCA, then the elapsed time within a 18-s segment was decoded. The width of the classification time bin was 1 s. **c.** Examples of confusion matrix of M2 and PPC. **d.** Comparison of decoding accuracy between M2 and PPC (Wilcoxon signed rank test, p=0.84). **e.** The decoding accuracies of M2 population, PPC populations, and mixed population of M2 and PPC were plotted against the dimension of the cPCA. Regardless of the dimensions, the decoding accuracies were similar. **f.** Comparison of the dimensions with the highest decoding accuracy between M2 and PPC (Wilcoxon signed rank test, p=0.25). **g.** Examples of the decoding results from M2 and PPC population. Black shade indicates the time of both M2 and PPC was correct. Yellow and green bars indicate independent and coherent error, respectively. **h.** Joint distribution of decoding errors relative to actual time in M2 and PPC. The right panel shows the category of errors. **i.** The rate of difference in decoded time when decoded results of both M2 and PPC showed error was plotted. The rate had a significant peak when the offset was 0 (p<0.001, bootstrap test). **j.** Comparison of consistent and independent error rate in each mouse (Wilcoxon signed rank test, p=0.03). **k.** Both consistent and independent error rates in each session were significantly greater than those of shuffled data (p<0.001, bootstrap test).

If M2 and PPC encoded time independently, mixing neurons from the two regions should improve decoding performance (Averbeck, Latham, and Pouget 2006). We therefore sampled equal numbers of neurons from M2 and PPC, and applied dimensionality reduction, and decoded time. Mixing neurons did not surpass the performance of M2 or PPC alone (**Fig. 3e**, black line), suggesting that the temporal representations in the two areas are not independent but mutually related.

### Two types of errors in the temporal representations of M2 and PPC

To assess cooperative computation across the fronto-parietal network, we next asked whether decoding errors arose independently or in a coordinated manner. Example trials are shown in **Fig. 3g**: in some cases, one area decoded correctly while the other erred (yellow), whereas in other cases both areas mis-decoded by the same amount (green). We refer to these as independent and coherent errors, respectively. The joint distribution of decoding errors displayed a cross and a diagonal band (**Fig. 3h**): the cross reflecting independent errors in which either M2 or PPC preserved the correct elapsed time, and the diagonal indicating coherent errors in which both areas deviated by the same interval. Quantifying trials in which both areas erred revealed a significant over-representation of cases where their timing errors differed by less than 1 s (**Fig. 3i**). Across sessions, independent errors were consistently more frequent than coherent errors (**Fig. 3j**; Wilcoxon signed-rank test, p = 0.03). Remarkably, the incidence of each error type varied little across animals (independent errors, 0.38 ± 0.013; coherent errors, 0.17 ± 0.010), yielding a ratio of 2.2 ± 0.14. In every animal both error types occurred at rates significantly higher than those in shuffled data (**Fig. 3k**). These findings indicate that while M2 and PPC generally maintain coherent temporal coding, they can also operate independently, and the balance between coherence and independence is tightly conserved across animals.

### CARP revealed the information carried by the shared subspaces

We hypothesized that coordination of temporal representations is achieved through corticocortical communication between M2 and PPC. To test this, we applied canonical-correlation analysis (CCA) to the simultaneously recorded population activities. CCA linearly transforms each population into an N-dimensional time series so that corresponding dimensions are maximally correlated (**Extended Data** Fig. 5). To determine what information each shared dynamic encodes, we developed **Canonical-component Ablation–based Representation Probing (CARP; Fig. 4a)**. CARP first establishes a baseline by applying CCA and decoding either elapsed time or lick rate from the full canonical components. Then, the N-th component is removed before decoding, and the new decoding accuracy is subtracted from the baseline; this difference quantifies the contribution of that component. Applying CARP to time decoding and lick-rate decoding revealed a striking dissociation. Time information did not depend most on the first (maximally shared) component but peaked at the fifth, whereas lick rate was encoded predominantly in the first (**Fig. 4b,c**). Moreover, temporal information was supported by nearly equal contributions from components up to about sixth, indicating a broadly distributed, high-dimensional code. By contrast, lick rate relied mainly on the first two components, implying a low-dimensional representation. Thus, the primary information that M2 and PPC share is dominated by behavioural signals, while temporal information is embedded more diffusely. We conclude that coordination of temporal representations across the fronto-parietal network is constrained by shared behavioural signals and is achieved through correspondences among high-dimensional, distributed population codes.

**Figure 4.**
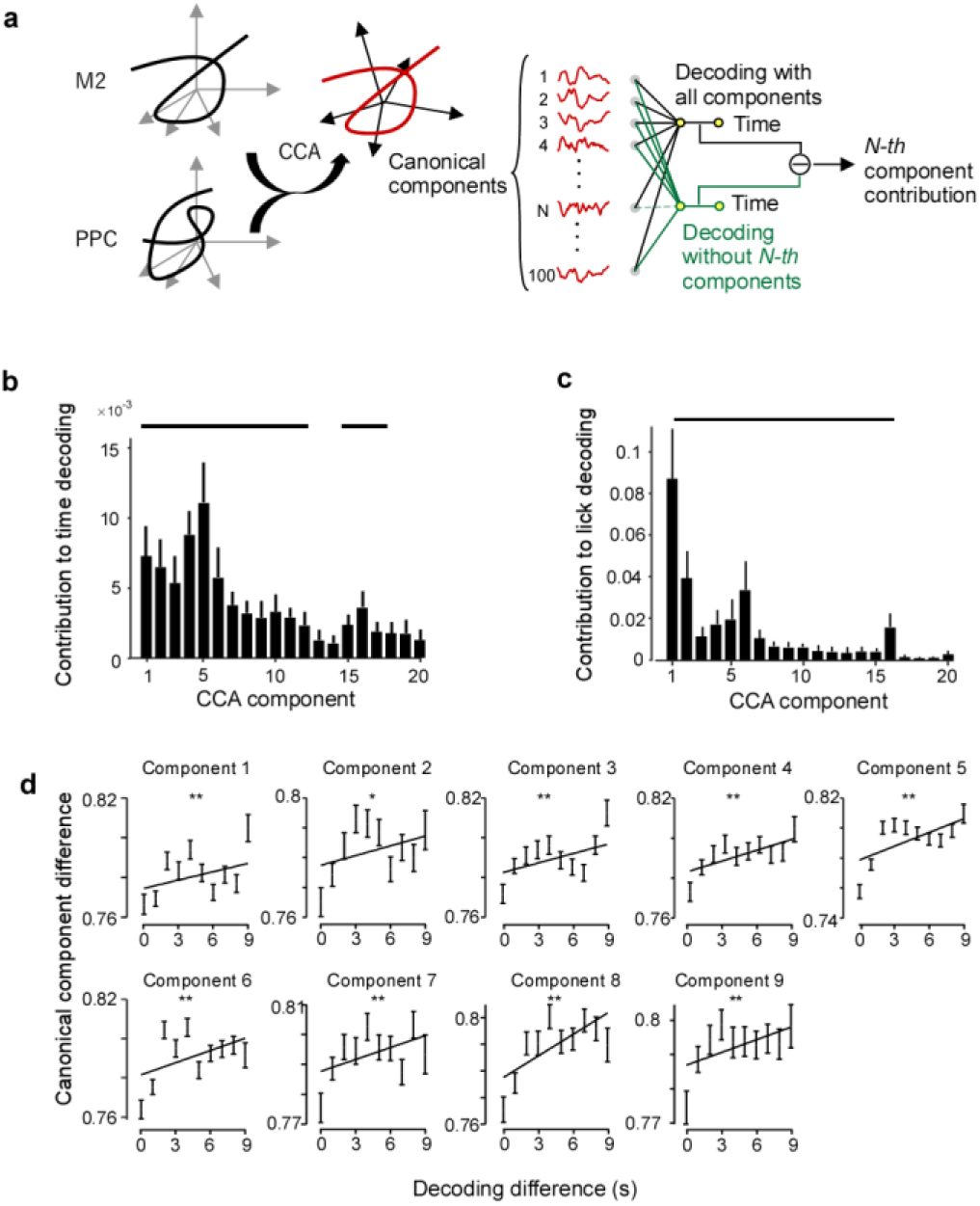
Shared M2–PPC dynamics encode time and behaviour. a. Schematic of CARP (Canonical-component Ablation–based Representation Probing). Common dynamics between M2 and PPC were extracted with CCA (red). The contribution of each component was assessed by deleting it from the full M2 activity and quantifying the resulting reduction in decoding accuracy. **b, c.** CARP results for decoding elapsed time (**b**) and lick rate (**c**). Bars denote components whose contribution was significantly larger than 0 (signed-rank test, *p* < 0.05, no correction). For each CCA component (1-9), the absolute difference between M2 and PPC scores was plotted against the absolute difference in the times decoded from the two areas (** *p*<0.01, * *p*<0.05, Spearman’s correlation test).

### Relationship between temporal coherence and corticocortical communication

To examine how corticocortical communication of each CCA component relates to the coherence of temporal representations, we compared the difference between canonical scores of M2 and PPC with the difference between their decoded times. Time bins in which the temporal estimates were more similar showed correspondingly smaller differences in canonical scores, and this relationship persisted up to 9-th component (**Fig. 4d**). These results qualitatively mirror the findings with CARP and strongly support the view that coherence of temporal coding across M2 and PPC is sustained by distributed, multidimensional shared dynamics between the two areas.

### Twin-RNN model emulating the dynamics of M2 and PPC

Analysis of the experimental data revealed two distinctive features of the fronto-parietal system. First, the two circuits maintain an intermediate balance between coherence and independence in their temporal codes: they neither move in perfect synchrony nor function as fully autonomous timers. Because the task demands no specific neural activity or overt behavior, this emergent balance is likely an intrinsic property of the cortex. Second, M2 and PPC share behavioural information primarily along high-variance population dimensions, whereas shared temporal information is dispersed across many lower-variance dimensions. The circuit mechanisms that support these dual properties and their mutual relationship remain unclear. To address this issue, we constructed a computational model in which each area is represented by a RNN (**Fig. 5a**), a framework well suited for probing continuous-time neural dynamics. The twin-RNN model comprises two networks, RNN-A and RNN-B, each containing 80 % excitatory and 20 % inhibitory units. The two networks are sparsely interconnected, with each synapse present at probability *p* = 0.001 (see **Methods**).

**Figure 5.**
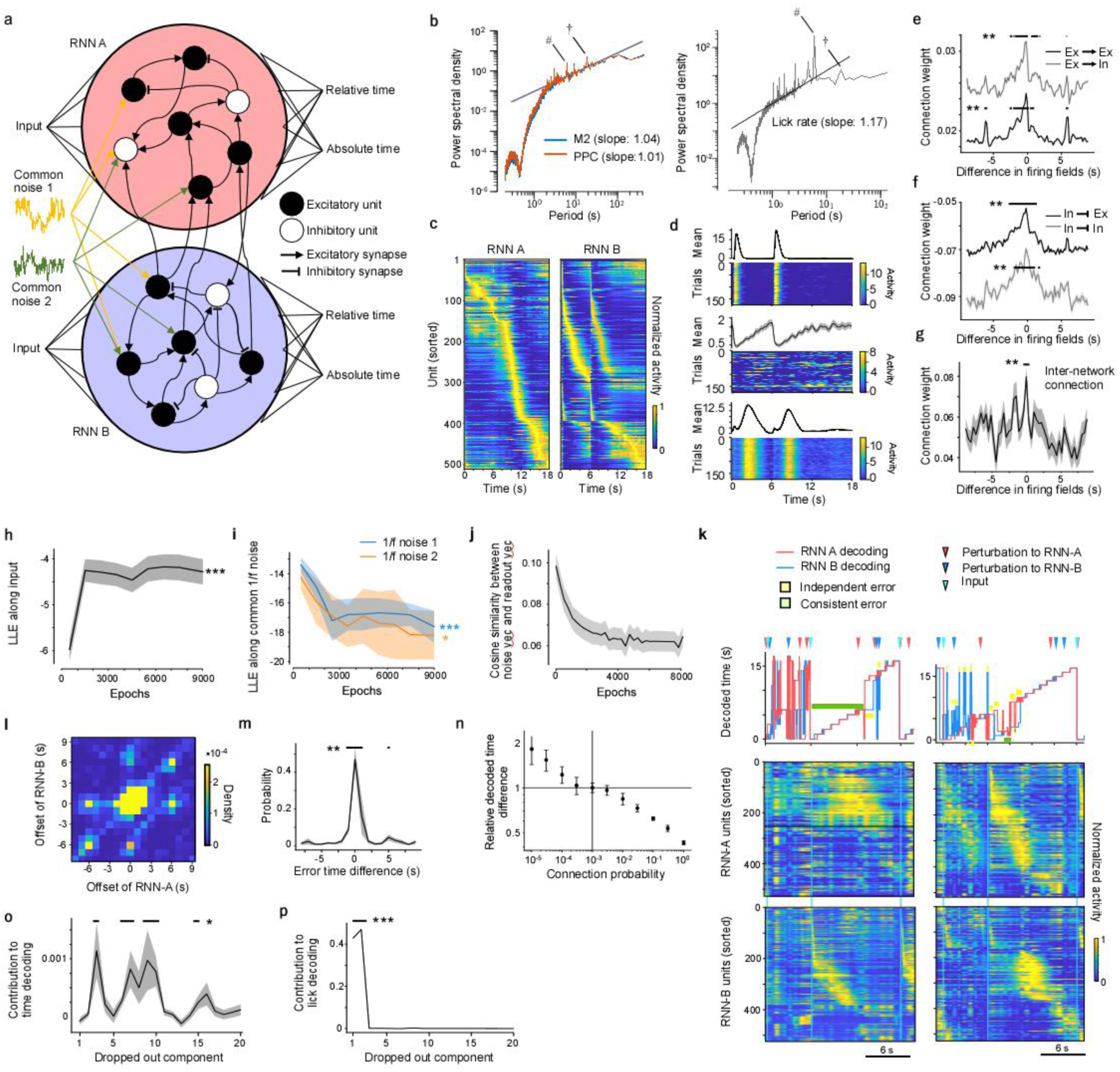
Twin-RNN model recapitulates key fronto-parietal dynamics. a. Schematic of the twin-RNN architecture. Each network (RNN-A, RNN-B) comprises 80 % excitatory and 20 % inhibitory units. Inter RNN projections are excitatory only. **b.** Power-spectral analysis of the first canonical component of the experimental data (left) and lick rate (right). The prominent 1/f component motivated injection of shared 1/f noise into the model. Lick rate showed a similar power-law statistics. # denotes peak at 6 s and †18 s. **c.** Trial-averaged activity of the trained twin-RNN units, sorted by peak time across 18-s interval. **d.** Representative task-related unit responses that emerged after training. **e,f.** Learned synaptic weights plotted against the temporal offset between the peak firing time (firing fields) of pre- and postsynaptic units, shown separately for excitatory (**e**) and inhibitory (**f**) presynaptic neurons. g. Inter-RNN connection weights plotted against the temporal offset between the firing fields of pre- and postsynaptic units. h. LLE along the input-vector direction over the course of learning. (*r* = 0.49, ***p = 1.7 × 10⁻^7^, Peason’s correlation test) i. LLE along the two 1/f-noise vectors over the course of learning. (1/f noise 1: *r* = -0.59, ***p = 2.6 × 10⁻^4^, 1/f noise 2: *r* = -0.22, *p = 0.026, Pearson’s correlation test) j. Absolute value of cosine similarity between the two 1/f-noise vectors and the readout vector over the course of learning. (*r* = -0.11, p = 1.5 × 10⁻^9^, Peason’s correlation test) k. Two examples of unit responses (bottom) and corresponding decoder outputs (top) following a transient perturbation to the RNNs. RNN units were sorted by the peak timing of trial averaged activity. l. Joint distribution of decoded-time offsets produced by RNN-A and RNN-B. m. The rate of difference in decoded time when both RNN-A and RNN-B showed error was plotted. The rate had a significant peak when the offset was 0 (p<0.001, bootstrap test). n. Using the default connection probability of *pc* = 10^−3^ as the baseline, we plotted the relative decoded time differences of RNN-A and RNN-B as a function of *pc*. o. CARP analysis on temporal information up to 20 Canonical components. Bars denote significant contribution (p<0.05 Wilcoxon signed rank test vs 0, no correction). p. CARP analysis on the shared noise of up to 20 CCA components. The first 2 components were significantly larger than the rest (p= 4.4 × 10⁻^5^, Wilcoxon signed-rank test).

To capture the shared, behaviour-linked fluctuations in M2 and PPC, we focused on statistical properties of spontaneous mouse movements and neural activity. It has been shown that behaviours, including licking, and neural activity follow scale-free statistics (Jones et al. 2023). Consistent with this, a spectral analysis of both lick rate and first canonical score of neural activity displayed a power-law profile (**Fig. 5b**). Since the latter decayed proportionally to the inverse of frequency (1/f), we injected 1/f noise into both RNNs. Because this noise is low- dimensional yet high-variance, it is strongly shared between the networks. Thus, the behaviour-related shared fluctuations observed in the experimental data were successfully incorporated into the model.

To reproduce in the twin-RNN model the behavioral capabilities observed in vivo, we trained the network to report both absolute and relative elapsed time in an alternating-interval (AI) task. After learning, RNN-A and RNN-B performed the task with high accuracy, confirming that the imposed synaptic architecture can support precise temporal computations. Inspection of single-unit activity revealed a rich repertoire of temporal response patterns. Alongside neurons whose profiles closely resembled the neurons identified experimentally, many additional motifs emerged (**Fig. 5c,d; Extended Data** Fig. 6). At the population level we quantified sequential organization using the sequentiality index and found that the model driven by the shared 1/f noise reproduced experimental values more faithfully than an otherwise identical model lacking this noise source (**Extended Data** Fig. 3a). Taken together, these results demonstrate that the twin-RNN receiving common 1/f fluctuations can not only master the AI task but also recapitulate the key dynamical signatures of M2 and PPC.

### Synaptic organization of learned twin-RNN

To understand what properties of the network made it possible to reproduce the dynamics, we first analysed the synaptic matrix after training. Among excitatory neurons, connection strength increased monotonically as the temporal distance between the pre- and postsynaptic firing fields decreased (**Fig. 5e**). In contrast, inhibitory connections became less negative when firing fields were nearby, indicating weaker inhibition between neurons that are active at similar times (**Fig. 5f**). These patterns imply that the slow population drift underlying the temporal code is generated by local excitation among neurons with similar firing times and by inhibition that preferentially suppresses units with dissimilar firing fields. Importantly, the same relationship held for the sparse inter-areal projections: the closer the firing fields of two neurons, the stronger their cross-area excitatory weight (**Fig. 5g**). This organization mirrors the structure inferred from the experimental data (**Extended Data** Fig. 3b-d).

Next, we investigated how learning reshaped the network’s responses to its inputs and to the shared 1/f noise. We quantified these changes by measuring how quickly the network returned to its attractor after being perturbed in specific directions, using the local Lyapunov exponent (**LLE**) (see **Supplementary Information**). The LLE evaluates the exponent λ that governs whether an error introduced along a given direction grows or shrinks exponentially: λ > 0 denotes exponential divergence, whereas λ < 0 denotes exponential decay. Thus, λ provides a direct measure of expansion or convergence speed. During training, the LLE along the input vector increased toward zero (**Fig. 5h**), while the LLE along the noise vector decreased (**Fig. 5i**). These trends show that the network learned to preserve the influence of inputs yet rapidly dampen the influence of noise. Consistently, the absolute value cosine similarity between the readout vector and the common 1/f-noise vector declined over training (**Fig. 5j**), showing that the network’s output became progressively orthogonal to the noise dimension and therefore more noise-resistant. Thus, training the twin-RNN induced at least four coordinated changes: synaptic weights organized neurons into temporally ordered firing sequences, the network learns to retain input information, noise was actively cancelled, and the readouts became orthogonal to the noise.

### Sparse inter-RNN connection and common 1/f noise reproduce experimental data

Next, we asked whether the twin-RNN reproduces the balance of independent and coherent errors observed in M2 and PPC. To test this, we injected brief perturbations to induce decoding errors (**Fig. 5k**). The model generated both independent and coherent errors: the joint distribution of decoded-time offsets from RNN-A and RNN-B contained a prominent cross (independent errors) and a diagonal ridge (coherent errors), exactly as in vivo (**Fig. 5l,m**). Thus, with a connection probability of p = 0.001, the twin-RNN retains, like the biological circuit, a simultaneous capacity for independence and consistency. We also confirmed that the decoding consistency increased with connection probability (**Fig. 5n**).

Next, we examined the shared network dynamics using CARP. Time decoding depended little on the first two canonical components and was instead supported broadly across higher components, confirming a high-dimensional, distributed code (**Fig. 5o**). By contrast, CARP with 1/f noise revealed that the common noise input was almost entirely included in the first and second components (**Fig. 5p**). Thus, the twin-RNN model faithfully captured the characteristic shared dynamics between M2 and PPC (**Fig.4 b,c**). These results show that even when the largest shared dimensions are dominated by common noise input that is irrelevant to the task, the two sparsely connected networks can align their high-dimensional temporal codes through the structured corticocortical synapses.

### Connection probability and perturbation direction determine inter-RNN coherence

To dissect how coherent and independent errors arise, we examined two factors: the inter-areal connection probability (*pc*) and the direction of external perturbations applied to the twin- RNN. We first analysed two extreme cases. When inter-areal coupling was absent (*pc* = 0), perturbations produced almost exclusively independent errors; conversely, with all possible cross-area synapses present (*pc* = 1), coherent errors predominated (**Fig. 6a**), indicating that cross-area excitation promotes coordination. Varying *pc* over 10⁻⁵–10⁻¹ showed that the rates of both error types decreased monotonically as coupling strengthened (**Fig. 6b**), yet their ratio (independent / coherent) declined steadily. Thus, inter-RNN coupling contributes chiefly by suppressing independent errors, thereby biasing the network toward coherent errors. These results imply that the consistency between M2 and PPC can be preserved by sparse corticocortical connections. At the same time, those connections stabilize the system as a whole and lower the overall error rate by making the temporal representation more robust.

**Figure 6.**
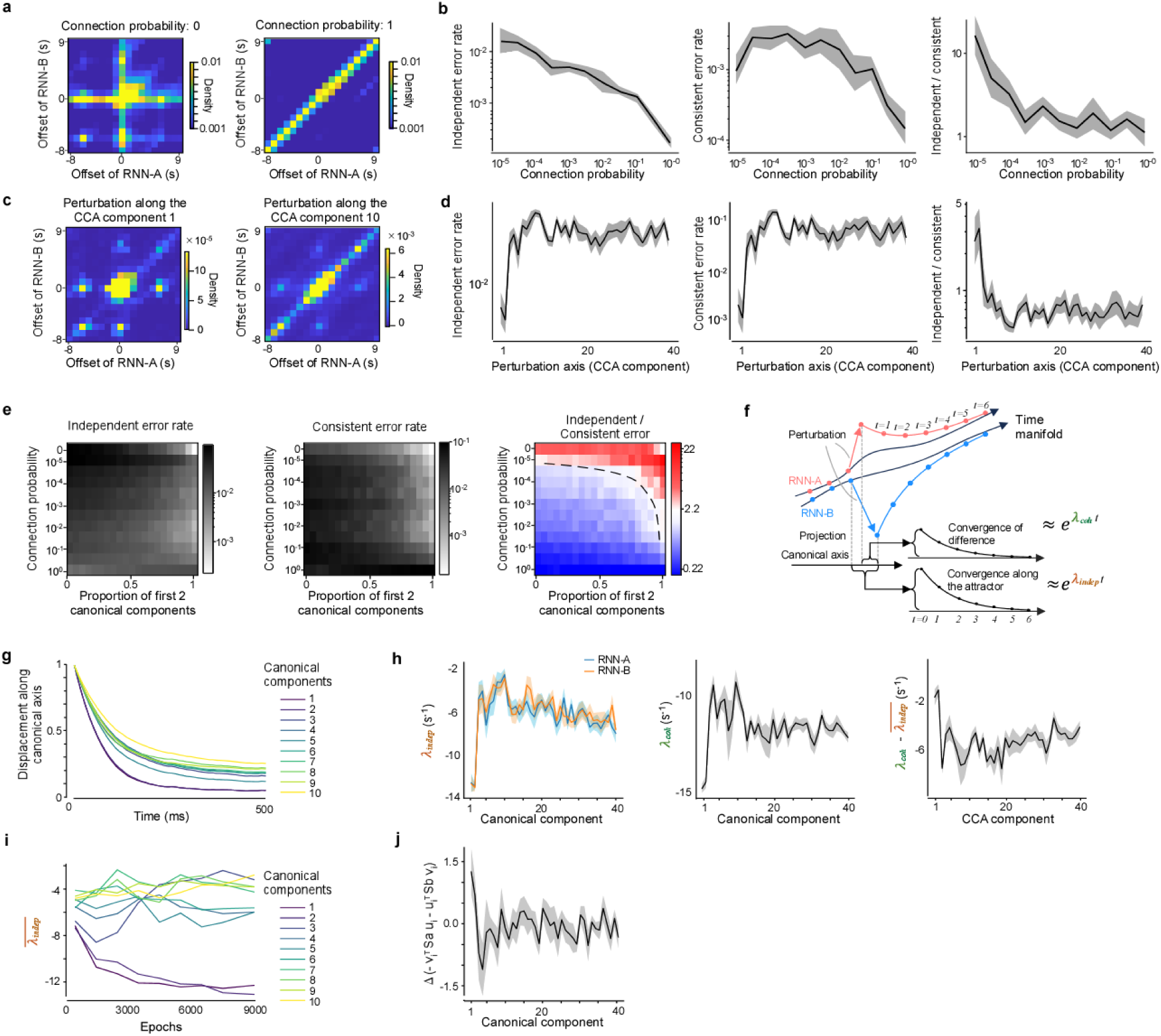
**| Coupling strength and perturbation axis jointly shape error independence and coherence in twin RNN** a. Joint distribution of instantaneous scalar errors in RNN-A and RNN-B when the inter-network connection probability is 0 (left) and 1 (right). b. Fraction of *independent errors* (**left**), coherent errors (**middle**), and ratio of independent to coherent errors (**right**) as a function of the inter-network connection probability *p*. c. Joint error distributions when a brief perturbation is applied to the 2nd canonical component (**left**) or the 10th component (**right**) of the shared state space. d. Fraction of independent errors (**left**), coherent errors (**middle**), and ratio of independent to coherent errors (**right**) as a function of the perturbed Canonical component index. e. Heat-map of the ratio in three variables in **b** and **d** plotted over the two-dimensional parameter space defined by connection probability (*p*, y-axis) and proportion of first and second canonical component (x-axis) within the perturbation noise. f. Schematic of the DLIC analysis. The **independent LLE** λindep quantifies the speed at which each RNN returns to its own attractor after a perturbation, whereas the **coherent LLE** λcoh measures the speed at which the two networks converge toward the corresponding states. g. Time course of the canonical score difference between RNN A and B after a displacement along each canonical component. Each line represents the result of perturbation along a specific canonical component. Displacements are normalized to be 1 at time 0. h. λindep (**left**), λcoh (**middle**) and their difference Δ𝜆 = 𝜆_𝑐𝑜ℎ_ − 𝜆̅_𝑖̅𝑛̅̅𝑑̅𝑒̅̅𝑝̅_ (**right**) plotted as functions of the perturbed canonical component. i. Training-induced changes in the local Lyapunov exponent (LLE) measured along the top 10 canonical components. j. Difference between approximate Δλ, −(𝒗^𝑇^𝑆_𝑎_𝒖_𝑖_ + 𝒖^𝑇^𝑆_𝑏_𝒗_𝑖_)at epoch 1 and epoch 8000, shown for canonical components n = 1–40. Negative values denote a decrease over training, positive values an increase through training. Components 1 and 2 exhibited significantly larger increments than the remaining components (p=4.9 × 10⁻^3^, Wilcoxon rank-sum test)

Next, to investigate the effect of direction of external perturbations applied to the twin-RNN, we delivered brief perturbations aligned with individual canonical components of the shared state space, with *pc* reset to the default 0.001 (**Fig. 6c,d**). Perturbation along the first two canonical components, dominated by common 1/*f* noise, favoured independent errors. In contrast, perturbations along canonical components 3–10, which carry the distributed temporal code, mainly induced coherent errors. These results show that error generation is strongly structured by perturbation direction: displacements along the shared temporal manifold yield coordinated shifts in both RNNs, whereas displacements along noise- dominated axes decouple their states. In the experimental data, the ratio of independent to coherent errors was remarkably stable across animals (∼ 2). To identify the conditions that meet these experimental results, we mapped the error-type ratio across the two-variable space defined by *pc* (10⁻^5^–10⁻²) and the proportion of the first 2 canonical components within perturbation noise (**Fig. 6e**). The target ratio of 2 emerged over a limited parameter space (dashed line in **Fig. 6e right**), demonstrating that the experimentally observed balance can be achieved by jointly tuning fronto-parietal connectivity and the directionality of perturbations rather than by connectivity alone. Together, these findings suggest that cortical circuits tune corticocortical projection patterns and the directions of externally driven perturbations to maintain the flexibility to switch the computational modes.

DLIC reveals how perturbations along the common-noise axes preferentially induce independent errors

To understand why perturbation along the highly shared-noise subspace gives rise to independent, rather than coherent, timing errors, we quantified two variables: stability of the attractor of each RNN, and speed of re-synchronization of RNNs. First, for each network we computed the LLE along each canonical component (**λindep**, **Fig. 6f,g**). −λindep quantifies how rapidly each RNN returns to (or leaves from) its attractor along a specific canonical component. The faster each network individually returns to its own attractor, the more likely it is to remain in the independent mode. All components exhibited negative λindep values (**Fig. 6h**), confirming that the system is ultimately stable in every direction. Notably, the first two CCA dimensions, dominated by the common 1/*f* noise, showed the most negative λindep values, indicating the fastest local convergence, whereas the time-coding dimensions converged more slowly. During the learning process, λindep decreased selectively in the first two components (**Fig. 6i**). These are consistent with the idea that the network during training learned to compensate for the activity changes along the noisy dimension while it learned to maintain the temporal information. We next measured the LLE of the difference between their CCA projections (**λcoh**, **Fig. 6f**). −λcoh quantifies how rapidly the twin-RNN re-synchronizes (or de-synchronizes) after perturbation. The faster the re-synchronization, the more likely the system is to enter the coherent mode. Again, all components exhibited negative λcoh values, confirming that two RNNs ultimately converge (**Fig. 6h**). The noise-dominated dimensions, the first two CCA dimensions, displayed the fastest re-synchronization, whereas the time-related dimensions were slower.

The large negative values of λcoh observed in the first two canonical components could arise either because two systems truly converge towards each other or simply because both collapse rapidly along the canonical components, thereby making their difference collapse quickly. To properly evaluate the synchrony the systems, we calculated a **D**ual **L**LE-based **I**ndependence vs **C**oherence (**DLIC**) index, defined asΔ𝜆 = 𝜆_𝑐𝑜ℎ_ − 𝜆̅_𝑖̅𝑛̅̅𝑑̅𝑒̅̅𝑝̅_, where upper bar denotes mean of RNN A and B. This quantifies the convergence rate between the two systems relative to their individual convergence rates. A large DLIC favours independence, whereas a small value favours coherence. The DLIC indexes of the first and second components were larger than those of time-coding dimensions (**Fig. 6h**). These results indicate that perturbations along the time-related axes quickly restore coherence between the two networks, whereas perturbations along the common-noise axes often let each RNN stabilize in its own attractor before re-synchronizing.

### DLIC is governed by sparse corticocortical connectivity and perturbation direction

We analytically solved DLIC (see **Supplementary Information**) and found that the index is determined by the alignment between inter-network connections and the perturbation directions applied to the two regions: strong alignment reduces the DLIC value, whereas weak or opposite (anti-parallel) alignment increases it. During training, the twin RNN gradually adjusted its inter-network weights such that their alignment with the first or second canonical component, the common-noise axis, declined progressively compared to other components (**Fig. 6j**). This suggests that the two RNNs learned to mutually suppress the influence of time-independent common noise. By contrast, for the time-coding dimensions (component 3 to 10), alignment changed little during training, which resulted in small DLIC values. These results replicated numerical observations. Thus, cortical computational modes are shaped by the consistency between perturbation directions and cortico-cortical connectivity, and that the network learns to organize this consistency so that specific perturbations reliably induce specific computational modes.

## Discussion

In this study, we developed a novel alternating-interval (AI) task that prompts the agent to update its internal state continuously while responding to external cues, and applied it to both behaving mice and RNNs. Using a Diesel2p mesoscope, we simultaneously imaged thousands of neurons in M2 and PPC at single-neuron resolution. The experiments revealed two key features. First, temporal codes in M2 and PPC can flexibly shift between coherent and independent modes. Second, their shared population dynamics are stratified: time information occupies a low-variance, high-dimensional subspace, whereas behavioral signals reside in a high-variance, low-dimensional subspace. To test whether these properties arise from minimal circuit ingredients, we built a twin-RNN model containing only sparse inter-network excitatory coupling (p = 10⁻³) and a common 1/f noise input. The model reproduced the experimental ratio of coherent-to-independent errors (1 : 2) and mirrored the same separation between temporal and behavioural subspaces. Further analysis showed how the network switches between computational modes. Taken together, our results suggest that sparse long-range connectivity, combined with large-scale shared noise, jointly governs the balance between independence and coherence in the fronto-parietal system.

To quantify the dynamics underlying different modes, we introduced a DLIC index, Δλ, by comparing (i) −λindep, the speed with which each area relaxes to its own attractor, and (ii) −λcoh, the speed with which the two networks return to the same states. Along the first two canonical components dominated by shared 1/ f noise, Δλ (λcoh − λindep) was close to zero. As a result, a perturbation allows each network to settle into its own attractor before the pair re-synchronizes. In contrast, along the time-coding axes (component 3–10) DLIC was negative, meaning the networks first realign their phases and only then stabilize, leading primarily to coherent errors. Thus, the global-noise axes act as “a source of independence,” whereas the sparse long-range excitatory projections serve as “sources of synchrony.” The experimental data suggests that these two influences are delicately balanced. Analysis based on the DLIC therefore offers a unified explanation of how local stability and corticocortical synchronization jointly generate both coordinated and autonomous temporal representations. Furthermore, our analytic solution of the DLIC revealed that Δλ mirrors the degree of alignment between inter-network connections and the perturbation directions, and that changes in these connections during learning shape the structure of Δλ. This framework is consistent with theoretical proposals that leverage controlled chaos to reconcile stability with diversity in temporal patterns (Laje and Buonomano 2013; Buonomano and Laje 2010), and it underscores the importance of multi- scale dynamical stability in achieving the coexistence of robustness and flexibility in biological time perception.

This study uncovered a functional role for the broadly shared, area-nonspecific cortical dynamics that have been observed across the neocortex recently (Stringer et al. 2019; Hira et al. 2024; Musall et al. 2019). Because such non-selective, behaviour-related activity may originate in the thalamus (Inácio et al. 2025), the largest shared component we detected is likely driven not by cortico-cortical projections but by a common thalamic input. The thalamus, in turn, receives modulatory signals from the basal ganglia, cerebellum, and superior colliculus as well as sensory afferents (Yoshida et al. 2024; Sugino et al. 2024); hence subcortical circuits could dynamically adjust thalamic drive, thereby switching global cortical operating modes. On the other hand, corticocortical functional connectivity can be dynamically modified by neuromodulators (Slater et al. 2022). Thus, by modulating the weight of common thalamic inputs or altering neuromodulator levels, the brain could accelerate area-specific processing or promote robust, coherent processing across areas. Amplifying shared noise is predicted to strengthen cortico-cortical correlations while paradoxically making information processing within each cortical area more autonomous. This leads to a testable hypothesis: boosting non-specific input or appropriately altering neuromodulators should increase inter-areal correlations, while simultaneously enhancing local computational independence.

The temporal coding mechanism uncovered here offers an instructive contrast to the continuous attractors that support spatial representations around the hippocampus. In the MEC, multiple grid modules normally maintain a coherent place code, though they can transiently decouple when the animal first encounters a novel environment (Stensola et al. 2012; Waaga et al. 2022). Because a spatial map must remain anchored to an absolute reference frame unless the surroundings change abruptly, coherence is favoured and strong inter-module coupling restricts the system’s degrees of freedom. Time, by contrast, lacks a natural zero point; ecologically, the brain must flexibly track the elapsed intervals from many different events. Consistent with this need for autonomy, independent errors were common in our experiments, and our modelling suggests that such independence is promoted by sparse long-range connection together with large shared noise. Nevertheless, spatial and temporal systems can be unified within a single scheme that comprises three elements: redundant copies of a continuous attractor, adjustable coupling among the copies, and substantial shared noise—any of which can lock or unlock relationships across brain areas. The analysis methods, CARP and DLIC, therefore provide a general dynamical lens that spans both spatial and temporal coding. Comparing space and time within this framework should advance a system-level understanding of brain-wide computation.

## Materials and methods

### Animal

All experiments were approved by the Animal Research Ethics Committee of the Institutional Animal Care and Use Committee of Tokyo Institute of Science (A2024-060C2) and were carried out in accordance with the Fundamental Guidelines for Proper Conduct of Animal Experiment and Related Activities in Academic Research Institutions (Ministry of Education, Culture, Sports, Science and Technology of Japan). A C57BL/6Jcl mouse and 4 Ai162(TIT2L- GC6s-ICL-tTA2) (Daigle et al. 2018) mice were used. These mice were kept under an inverted light schedule (lights off at 12 AM; lights on at 12 PM) in their home cages to adapt to experimental surroundings.

### Neonatal AAV injection

The injection pipette was fabricated from 10-mm-long borosilicate glass capillaries (outer diameter = 1.00 mm, inner diameter = 0.75 mm; no filament, B100-75-10, Sutter Instrument). Capillaries were pulled on a micropipette puller (P-97, Sutter Instrument) to yield tips with an outer diameter of ∼50 µm and an inner diameter of ∼30 µm. The pulled pipette was connected to silicone tubing, which in turn was attached to a 10-µl microliter syringe (801 RN, Hamilton Company). Viral injections were carried out on postnatal day 1 in three pups. Each pup was anaesthetized by hypothermia on an ice bed and secured in a custom neonatal head holder designed by Mr Ooishi (Narishige). The holder’s arms were set at a 20° incline, gently grasped the neck 7–9 mm apart, and rotated the head ∼40° from the midline to improve access to the injection site. A pulled glass micropipette was advanced perpendicularly through the skull and then lowered 250–300 µm. A total of 4 µl of AAV-DJ-syn-jGCaMP8s (1.16 × 10^13^ GC/ml) was delivered at 4 µl/min with a syringe pump (Legato 100; KD Scientific). Both hemispheres were injected in two mice, and only the right hemisphere in one mouse. Pups were placed in a heated recovery chamber and returned to the dam once normal activity resumed. The entire procedure required <10 min per pup.

### Local injection of AAV

For two mice, 0.2 µl of AAV9-syn-jGCaMP8s-WPRE (1.25 × 10^13^ GC/ml) was pressure - injected into bilateral M2 and PPC at a depth of 500 µm from the pia, using a syringe pump (Legato 100; KD Scientific) set to 0.02 µl/min. Injections were delivered through borosilicate glass capillaries without filament (B100-75-10; Sutter Instrument), identical to those used for the neonatal AAV injections. Capillaries were pulled on a micropipette puller (P-97; Sutter Instrument) and the tips were bevelled to 45° with a Narishige grinder, yielding a tip outer diameter of ∼40 µm and inner diameter of ∼30 µm.

### Behaviour training

A custom training system was used to train head-fixed mice. Head plate fixature and mouse body holder were designed with CAD and fabricated with a 3D printer. Lick detection device (Slotnick 2009) and water delivery pump were controlled by a single custom-written script (LabVIEW 2019, National Instruments). Throughout training a continuous 8-kHz pure tone was played to mimic the acoustic noise of the resonant scanner used during two-photon imaging. Following recovery from head-plate implantation and large cranial-window surgery, mice were water-restricted to 1 ml/day until their body mass reached 80–90 % of the pre-restriction weight. Head-fixed mice were then given 2 μl of water rewards at a 6-s fixed time interval. Each session typically lasted for 200 to 300 trials. We observed licking just before the reward delivery 3-6 days after the initiation of training. For each session, average lick rate at 0.2 s before the reward and the minimum of average lick rate between 1.5-4 s after the reward were monitored. When there were at least two sessions where the difference between former and the latter exceeded 3 Hz, the mouse was considered to have mastered 6s fixed interval task. The mouse then moved onto the alternating interval task, where it was given water rewards at 6-s and 12-s periods alternately. The mouse was considered to have mastered the task when the difference between the average lick rate at 11.8 s and 6 s in 12-s period exceeded 1 for more than five sessions. After the animal had completed at least one training session under the two-photon microscope, imaging sessions were initiated.

### Head plate surgery

Mice were anesthetized with isoflurane (1-1.5%) and the body temperature was monitored with a rectal thermometer (RET-3 Rectal Probe for Mice; Physitemp Instruments). Eye ointment (Tarivid ophthalmic ointment 0.3%; Santen Pharmaceutical) was applied to keep them moist. Head plate surgery was conducted on mice over 8 weeks of age (Hira et al. 2013). Scalp overlaying the dorsal cortex was removed before detachment of peritoneum and cranial muscles from the skull. A custom head-fixing imaging chamber was mounted on the skull with a self-cure dental adhesive resin cement (Super bond, L-Type Radiopaque; Sun Medical) and luting cement (FUJI LUTE BC; GC Dental). The ample width of the head plate allowed the implantation of a 7 mm × 7 mm glass plug and subsequent two-photon imaging. The surface of the intact skull was then coated with clear acrylic dental resin (Super bond, Polymer clear; Sun Medical) and dental silicone (Dent Silicone-V Regular type; Shofu) to prevent drying. The mouse was administered 0.38 μl / body weight (g) of meloxicam subcutaneously to reduce inflammation and pain. The mouse was then returned to the cage to recover for at least three days.

### Cranial window implant

Three days after head plate surgery, we performed an ultra-large cranial window implant. The glass plug used to cover the dorsal cortex was composed of a 6 mm × 6 mm #3 cover glass and a 7 mm × 7 mm #0 cover glass, both custom-manufactured by Matsunami Glass Ind. The front two corners of each cover glass were diagonally cut, resulting in a hexagonal shape. Two coverslips were glued together using optical adhesive (Norland Optical Adhesive 63; Norland Products). After anaesthesia, 5 μl/body weight (g) of 20% mannitol solution was administered via intraperitoneal injection (Takahashi, et al., 2021) to reduce intracranial pressure, and 0.216 μg/body weight (g) of atropine sulphate was administered intraperitoneally (Goldey et al. 2014) to reduce saliva and mucus secretion. The mouse head plate was fixed after anaesthesia. Clear acrylic dental resin was removed using a dental drill followed by marking of bregma and stereotaxic coordinates. The skull along the outline of the cranial window as well as coronal and sagittal sutures was thinned with the drill. The frontal and parietal bones were removed in four separate pieces. The glass plug was then placed on the exposed cortex, enabling optic access to the entire dorsal cortex. The glass plug was attached to the bone with cyanoacrylate glue and secured using dental resin cement. Finally, the mouse was administered 10 μl / body weight (g) of mannitol intraperitoneally and 0.38μl / body weight (g) of meloxicam subcutaneously before returning to the cage to recover for five days.

### One-photon calcium imaging and sensory stimulation

Imaging was conducted in a dark environment with the mouse head-fixed to restrict movement. Blue LED light (470 nm; light source: M470F4, Thorlabs; excitation filter: FF01-469/35-25, Semrock) was delivered to the cortical surface from the upper right side of the brain at a 45° angle. Violet LED light (405 nm; light source: M405FP1, Thorlabs; excitation filter: FBH405- 10, Thorlabs) was delivered from the upper rear right side (−45° azimuth, 45° elevation). The two excitation lights were alternately applied to the cortical surface at a frequency of 20 Hz (24 ms per illumination). The LED driver (LEDD1B, Thorlabs) was controlled using a microcontroller (Teensy 4.0, PJRC) with custom code written in Arduino IDE 2.2.1 (Arduino). The intensity of the excitation light was optimized for each animal and session. Fluorescence signals from the brain were collected through an objective lens (NIKKOR 85 mm F2, Nikon), an imaging lens (NIKKOR 50 mm F1.4, Nikon), and an emission filter (FF01-525/39-25, Semrock), and recorded using a CMOS camera (Alvium USB 3.1, Allied Vision). Image acquisition was controlled with custom software written in LabVIEW 2020 (National Instruments). Imaging was performed at 504 × 504 pixels (with a pixel size of 14.1 μm × 14.1 μm), with 3 × 3 binning, an exposure time of 24 ms, and a frame rate of 40 Hz (20 Hz × 2), such that signals under blue and violet excitation were captured alternately, frame by frame. The focal plane was manually adjusted to the dorsal surface of the cortex.

In somatosensory stimulation experiments, awake head-fixed mice received vibratory stimuli to trunk, tail, or left/right whiskers). These vibratory stimuli were designed to evoke both cutaneous and proprioceptive somatosensory responses. The 50 Hz stimuli were delivered by attaching a lightweight sterile plastic pipette (EOG sterilized poly dropper, 2 ml, Asone) to the vibrating element of a speaker (MM-SPL2NU3, Sanwa Supply) and placing it in contact with the mouse body. Stimulation was controlled by custom LabVIEW 2020 software (National Instruments). Each stimulus lasted 750 ms, with an interstimulus interval of 7500 ms. In each session, approximately 220 stimuli were presented for ∼30 minutes. For data analysis, signals were averaged across all trials and represented as mean brightness values. For three mice, the location of peak response to each stimulation was mapped and aligned with the mouse brain atlas. For two mice, the dorsal cortex and the brain atlas (Allen Mouse Brain CCF) was aligned using stereotaxic coordinates identified during cranial window implant surgery. The anterior, anteromedial, and rostrolateral visual areas were defined as PPC (Hira et al., 2025).

### Two-photon calcium imaging

Two-photon imaging was performed with a custom-built mesoscope (Diesel2p (Yu et al. 2021)). Excitation was provided by a fixed-wavelength femtosecond fibre laser (ALCOR 920-2, 2W, 80 MHz, Spark Lasers, France). Group-delay dispersion (∼60,000 fs²) was pre-compensated inside the laser head by maximizing sample fluorescence. Laser power was regulated with a half-wave plate followed by a polarizing beam-splitter cube. The beam was then expanded two- or three-fold (×2 or ×3 beam expander, Thorlabs) to fill the objective’s back aperture. Beam steering employed a resonant scanner (CRS 8 kHz, Cambridge Technology) in series with two galvanometric mirrors (R6210H, Cambridge Technology), all driven by an analog output board (PCI-6722, National Instruments). Custom scan and tube lenses, a primary dichroic mirror, the objective, and the collection optics completed the optical path. Sample power was usually 80-100 mW and never exceeded 130 mW in any experiment. The imaging depth was at layer 2/3 (100-150 μm from pia). Fluorescence was routed through the dichroic mirror and a 700 nm short-pass emission filter (FESH0700, Thorlabs) and detected by a GaAsP photomultiplier tube (H10770PA-40, Hamamatsu). The signal was amplified (#59-179, Edmund Optics) and digitized at 12 bit, 10 MS/s (ATS-460, AlazarTech). A line-sync signal from the resonant scanner provided the master clock for all devices. Custom LabVIEW software (LabVIEW 2019, National Instruments) controlled acquisition. For mice expressing GCaMP broadly across the dorsal cortex, two adjacent tiles (4.25 × 1.5 mm each) were collected and stitched to yield a 4.25 × 3 mm field of view (FOV) at 1,500 × 1,168 pixels and 4.79 fps.

### Image processing

Image preprocessing was done using ImageJ (National Institutes of Health) and MATLAB (MathWorks, Natick, MA, USA) software. The bidirectional phase offset of the image was calculated, and sinusoidal distortions caused by the resonant scanner were corrected. The Turboreg plugin (Thévenaz, Ruttimann, and Unser 1998) on ImageJ was used for image registration. Using ImageJ macros, the images were divided into smaller blocks of 150×150 pixels, and registration based on translation was performed for each block. Subsequently, MATLAB was used to combine the drift-corrected images from each block. The xy offset of each block was linearly interpolated across all of the pixels to ensure consistency of the shift. The registered images were then analysed using Suite2p (Pachitariu et al. 2016) to determine the location of the regions of interest (ROIs). In addition to the automatically detected ROIs, we visually inspected the images and manually added any morphologically recognizable neurons. Conversely, any ROI whose mean fluorescence intensity was lower than the background was excluded from further analysis. The intensity of each ROI was calculated by subtracting mean values of pixels within 120 μm of the ROI excluding the other ROIs from the mean value of the ROI. For each ROI, the 8 percentile pixel values within 15 s of the time window were obtained, which was then smoothed using a Gaussian filter with standard deviation of 30 s. This was considered as the baseline of neural activity, and was subtracted from the original data to set the baseline to 0 (Hira et al. 2013). A Wiener filter with temporal length of 1.5 s was applied to reduce noise. Non-negative deconvolution with time constant of 0.87 s (fast-oopsi; (Vogelstein et al. 2010)) was performed. Finally, gaussian filter with standard deviation of 0.2 s was applied for all analyses, except for decoding analysis, where a standard deviation of 0.5 s was used. For analysis, the neural data was interpolated to have a 10 Hz signal rate.

### Single neuron analysis

ROIs whose distribution of the activity had skewness over 0.3 were defined as **“reconstructed neurons”**, which were included in the analysis (Hira et al. 2013). Reconstructed neurons were classified as “**absolute timing neurons”** or “**relative timing neurons”** if the time ratio of the time of the maximum activity during 12 s and 6 s were between 0.9 to 1.1, or 1.9 to 2.1, respectively. Task-related neurons whose activity peak was within 2 s after reward were defined as reward-related neurons, and these neurons were excluded from the absolute time neurons or relative time neurons.

### Population activity characterization

To quantify sequential firing activity of neurons, peak entropy, temporal sparsity, and sequentiality index (Zhou, Masmanidis, and Buonomano 2020) were calculated. One hundred neurons were selected randomly from M2 and PPC, and the data were averaged over trials. The indices were calculated as follows,

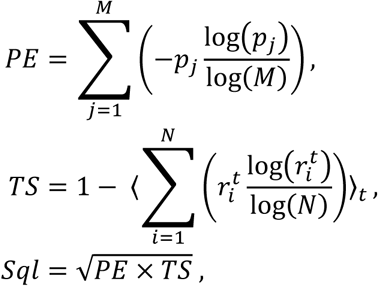

where M is the number of bins used to estimate the distribution of peak times (60 for 6-s period and 120 for 12-s period). 𝑝_𝑗_ is calculated as the proportion of units whose peak activity falls into time bin 𝑗, relative to the total number of units. 𝑟^𝑡^ indicates the activity of unit i at time bin 𝑡, normalized by the total activity across all units at that time. The notation 〈 〉_𝑡_ represents averaging over time, and 𝑁 denotes the total number of units.

𝑃𝐸 (peak entropy) quantifies how spread out the peak times are across all neurons in the population whereas 𝑇𝑆 measures how peaked or uneven the normalized activity is at each time point. The 𝑆𝑞𝐼 (sequentiality index) approaches 1 when neurons have peak times that are evenly distributed across the whole time period, such that at every moment, exactly one neuron is active and their temporal activity windows do not overlap.

The indices were calculated for 6-s period and 12-s period each, and the average of 100 iterations were analysed. When calculating sequentiality index of RNN result, the data was averaged per 10 samples to match the sampling rate of neural data (10 Hz).

### Noise correlation analysis

For each neuron pair, Pearson’s correlation coefficient between trial-to-trial fluctuations, defined as deviation from the mean activity, was calculated separately during the 6 s and 12 s intervals. The correlation was plotted against absolute difference in peak firing times, where peak times were taken from the mean activity in the alternate interval. Analyses were performed on three groups of pairs: within M2, within PPC, and cross-area (M2–PPC). To avoid edge-of-trial artifacts, any neuron whose firing peak occurred within 1 s before or after reward delivery was excluded.

### Encoding model

The encoding analysis was performed using ridge regression. The activity of each reconstructed neuron was fitted using lick rate and elapsed time within trial. Elapsed time was composed of 18 gaussian kernels where 𝑖th kernel had a peak at 𝑖th second in the trial. The standard deviation was 0.9 s. The data was intervals of 18 s each, consisting of 6 s interval and 12 s interval of the behavior task. These segments were then divided into 4 groups based on the modulo 4 of their segment order numbers. One group was used for testing while the remaining groups were used to fit regression models. The ridge parameter was set to 10^-4^ for all neurons.

R-squared was used to evaluate the model’s fit with the neural data. To evaluate the contribution of each predictive variable, permutation test was performed. Each variable was randomly shuffled in the time axis, and the regression was repeated 100 times. If the R² of the full model exceeded the 95th percentile of that variable’s shuffled R² distribution, the neuron was considered to be significantly associated with that predictor.

### Dimensionality reduction

Dimensionality of the neural data was reduced using a slightly modified version of contrastive principal component analysis (cPCA) (Abid et al. 2018) and canonical correlation analysis (CCA). For mathematical details, see **Supplementary information**. In brief, in our analysis, cPCA finds axis 𝑣 that maximize the intra-trial variance while simultaneously minimizing inter-trial variance. Parameter w determines the balance between the two. We changed the parameter w to find the optimal balance of within-trial variance and across-trial variance by comparing the decoding performance (**Extended Data** Fig. 4a,b). The decoding performance based on the cPCA was also compared with other dimensionality reduction methods, such as PCA, demixed PCA (Kobak et al. 2016), or Task-related component analysis (TRCA) (Tanaka, Katura, and Sato 2013) (**Extended Data** Fig. 4c). dPCA was done by dividing each trial into 1-second bins (18 pseudo-conditions) and, for each bin, computing its mean deviation from the overall trial average. Using this condition-dependent signal, condition-dependent encoding axes and its projection were calculated. TRCA seeks to find the axes that minimize average covariance within each single trial while maximizing the covariance of the trial-averaged responses.

### Neural manifold analysis

After dimensionality reduction with cPCA, the data was visualized using UMAP implementation in MATLAB. In figure 3a, neural data was transformed with cPCA (w = 0.6), and top 30 dimensions were used to visualize activity using UMAP (n_neighbors = 15, min_dist = 0.3). Comparison of neural manifold across range of cPCA variable w was done using neural data averaged per 5 trials (**Extended Data** Fig. 4a). The first 10 dimensions of the transformed data were subsequently transformed by UMAP (n_neighbors = 20, min_dist = 0.01) for visualization.

### Decoding

Decoding of temporal information from a population activity was done with a bagged decision tree algorithm (MATLAB function, *treebagger)*. The number of trees, a hyperparameter of TreeBagger, was fixed at 100. The neural data was segmented into intervals of 18 s each, consisting of 6 s interval and 12 s interval of the behaviour task. The classifier predicted the within trial time in integer values ranging from 1 to 18, i.e. 18 classes of each 1 s time window. To avoid the classifier from naively predicting based on temporally close training intervals, the trial segments were divided into 8 groups based on the modulo 8 of their segment order numbers. When decoding data at group i (ranging from 1 to 8), all data except for group i-1, i, and i+1 was used to train the classifier. The training data’s dimension was reduced using cPCA, PCA, dPCA, or TRCA, and the same coefficient was used to transform the testing data. The decoding analysis was done using dimensionality ranging from 2 to 300, and the optimum number of dimensions was identified for each mouse. Decoding accuracy was defined as a proportion of predictions that fell within 1 s of the real time. The cPCA parameter, w, was set to 0.6 for M2 and 0.5 for PPC, as these values yielded the highest average decoding accuracy.

### CARP analysis

Neural data from each area was reduced to 100 dimensions using PCA to prevent overfitting by CCA. Then the time series was transformed by the M2–PPC CCA projection. Elapsed time and lick rate were decoded, and the resulting accuracy was taken as the maximal decoding performance. Next, we removed the *N*-th CCA dimension (*N* = 1, 2, … , 100), repeated the decoding, and recorded the new accuracy; the reduction relative to the maximal decoding performance was interpreted as the contribution of that component. For elapsed time decoding, the data’s dimensionality was reduced using cPCA (w=0.6) and dimension with highest decoding accuracy was selected. For lick rate decoding, lick rate was obtained by counting licks in 0.4-s bins and was decoded with linear ridge regression. To prevent overfitting, an eight-fold cross-validation was used and performance was quantified by the coefficient of determination, 𝑟². The ridge regularization parameter was swept from 10⁻¹⁰ to 10¹⁰, and the value yielding the highest 𝑟² was adopted for all subsequent analyses. For every session, two independent, non-overlapping random samples of 100 neurons were selected from either M2 or PPC, and the entire analysis was repeated.

### Noise analysis

The neural data from M2 and PPC were first transformed by CCA. For each canonical component in M2 and PPC, Power Spectral Density (PSD) was calculated using *pwelch* function in MATLAB. PSD was plotted against period (s) in the log-log axis. PSD of period greater than 1.5 s was linearly fitted. The coefficient of the slope of the first canonical component was close to 1. This indicates that the slow baseline fluctuation was the 1/f (pink) noise. Lick rate was analysed in the same way, and the PSD from 0.8 s to 10 s period was linearly fitted.

### Single RNN models

A single-RNN model was derived from one branch of the twin-RNN architecture: a fully connected network of 512 units. For the alternating-interval task, the single RNN was trained on the same target output to the twin-RNN case. For the elapsed-time task, the network received two distinct input vectors. The inter-trial interval (ITI) was randomly drawn from 1.5– 4 s. Targets and learning rules were identical to those used for the twin-RNN. In the delay-matching-to-sample +time task, two input cues were presented with either a 6 s or 12 s delay. If the cue identities matched, the network was required to output both the relative and absolute 6-s interval; if they differed, it had to output the 12-s interval. An ITI of 1.5 – 4 s was imposed. Training procedures including loss functions, optimization, and stopping criteria were the same as for the twin-RNN model.

### Unusual input timing experiment

The *RandomForestClassifier* function from scikit-learn was used to classify activities of the RNN to either 6-s or 12-s period. The activities were dimensionally reduced to 100 dimensions using cPCA (w=0.6). The number of estimators was set to 100. For the 6-s period, we randomly applied input at unusual timing to the network. The activity of one second after the unusual input was decoded, and the number of times the decoder predicted 12-s period were counted to calculate the probability of transitioning from 6-s to 12-s period. For each unusual input timing, the experiment was repeated for 16 times, and the probability of transition was averaged. The probability was calculated similarly for the 12-s period. For the elapsed time task, inter trial intervals were removed from training datasets and the same analysis as the AI task was conducted using two different cues.

### Twin RNN models

The dynamics of the two RNNs, A and B, was described by the differential equations below.

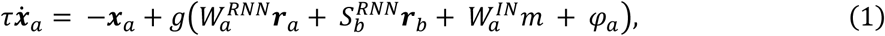

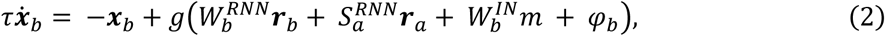

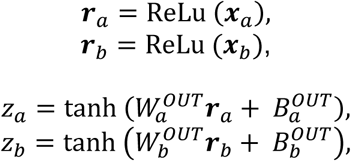

where 𝑥_𝑎_ and 𝑥_𝑏_ represented membrane potential and 𝑟_𝑎_ and 𝑟_𝑏_ represented firing rate of units in RNN A and B, respectively. Both membrane potential and firing rate were expressed as N by 1 vector, where N was the number of units set to 512. The firing rate was given by applying ReLU function on the membrane potential. The function, *g*, represents leaky ReLU with alpha=0.2. 𝑚 is the scalar input, and 𝜑 is Gaussian noise vector (*N*×*1*) with 0 mean and 0.08 variance. Hyperbolic tangent function was used for the activation function of the output 𝑧. The time constant 𝜏 was set to 100ms and the discretization time step was 10 ms. Following Dale’s law, 80% of the units were excitatory and 20% were inhibitory. Output connection weights from excitatory units were nonnegative whereas that of the inhibitory units were nonpositive. Connections between RNNs were constrained to be nonnegative, as is the case with cortico-cortical neurons. Connection strength between 2 RNNs was varied by setting connection probability (0 to 1) of the connection matrix Sa and Sb. In the default conditioning, the probability was set to 10^-3^ based on an anatomical study (Gămănuţ et al. 2018), a modelling study (Saiki-Ishikawa et al. 2025), and a theoretical study (Horvát et al. 2016). The maximum value of the synaptic weight was set to 2.

Two RNNs received a single cue corresponding reward or two different cues for 30 ms, three time-steps. The input weights underwent plasticity during training as the other synaptic weights. The input cue was given at 6 s and 12 s intervals alternately, analogous to the behaviour task. The output weight of each RNN for absolute and relative times was also plastic during training. Two outputs of each RNN were trained to fit one of which was a linear signal reflecting absolute time, and the other was a ramp-up signal in the form of quartic function which was proportional to the interval reflecting relative time from the input The maximum value of the input and output weights was set to 2.

Common noise was modelled as two independent 1/𝑓 processes chosen to match the experimental power spectrum for periods ≥ 1 s (frequencies ≤ 1 Hz). Each trace was rescaled so that its maximum and minimum values were +0.5 and –0.5, respectively. Then, the trace was re-scaled by a gain factor 𝜎 before multiplied by noise vector (*N*×*1*) and added to the RNN input vector 𝜑. The noise vector for the two components (noise vector 1 and noise vector 2) remained constant throughout all simulations.

### RNN training

The training of RNNs without shared noise input was done with batch size of 8, and each batch consisted of 12 trials for twin RNN models and 8 trials for single RNN models. The loss was calculated by taking the squared error of all 4 outputs with respect to 4 target outputs, consisting of relative time and absolute time from two RNNs, and taking the entire mean. In the case of a single RNN model, the number of targets was two. The model was trained by back propagation through time (BPTT) with Adam optimizer (learning rate = 5 × 10⁻⁵, clip value = 5 × 10⁻⁵ ). L2 regularization (coefficient = 0.1) was applied to the unit activities during training.

To reduce memory load during BPTT training, the time sequences were split into segments (12 for twin RNN models, 8 for single RNN models) using a stateful RNN in *tensorflow*, with parameters updated after processing each segment. Note that the synaptic plasticity was applied to all the synapses including within RNN synapses and inter RNN synapses regardless of the inhibitory and excitatory ones. The sign of the synaptic weight was 0 or positive for the excitatory synapses and 0 or negative for the inhibitory synapses. The output errors during the first trial were not included for calculating the loss since the model has no telling what interval comes first.

Training of the RNNs that received shared noise input was conducted with a batch size of 64, each batch containing 12 trials. A network previously trained without common noise served as the initial state. During the first 8,000 epochs the noise scaling factor σ was linearly ramped from 0 to 3; training then continued for another 2,000 epochs with σ fixed at 3. For analysis we selected, from this final interval, the model exhibiting the lowest loss. Optimization used the Adam algorithm (learning rate = 2.5 × 10⁻⁵, clip value = 2.5 × 10⁻⁵).

For each connection probability, the twin-RNNs were trained 10 times using different random seeds. Unless otherwise noted, all analyses were performed across the 10 models and the results were averaged.

### Weighted perturbations along CCA components

We split the CCA space into its first two components and the remaining 98 components and defined a perturbation vector 𝒗(𝑑, 𝑝) such that the ratio of the summed squared weights satisfies

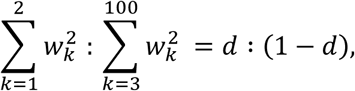

with ‖𝒗(𝑑, 𝑝)‖ = 1. Here 𝑑 ∊ [0,1] specifies the relative weight assigned to the top two CCA components. The unit vector 𝒗(𝑑, 𝑝) was then multiplied by a random scalar drawn from a normal distribution 𝑁(0, 18^2^) and added to the RNN input. For each combination of d and the inter-areal connection probability p, we quantified the frequencies of independent and coherent errors, as well as their ratio (**Fig. 6g**).

### DLIC (dual local Lyapunov exponent-based Independence vs Coherence) analysis

For detailed mathematical formalization, see **Supplementary information**. To assess the stability of the RNN, we used the concept of the local Lyapunov exponent (Waldner and Klages 2010)(Waldner and Klages 2012). Instantaneous stability of RNN A along 𝑖 th canonical direction, 𝑢_𝑖_ (normalized), is

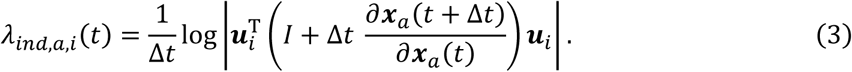

Similarly, for RNN B and its corresponding 𝑖th canonincal direction 𝑣_𝑖_, the LLE is

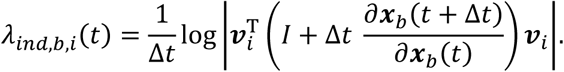

activity of RNN A and B at time t are denoted by 𝑥_𝑎_(𝑡) and 𝑥_𝑏_(𝑡), respectively. The parameter 𝜆_𝑖𝑛𝑑,𝑎,𝑖_(𝑡) and 𝜆_𝑖𝑛𝑑,𝑏,𝑖_(𝑡) represents the exponential rate of convergence or divergence along the attractor of each RNN at time t. Negative values indicate convergence (stability), and positive values indicate divergence (instability) of the trajectory projected to the chosen axis. The implementation for calculating local Lyapunov exponent was done on

TensorFlow. Using Eq. (3), local Lyapunov exponents along isolated canonical components at time 𝑡 were calculated via TensorFlow’s automatic differentiation. Isolated canonical components were obtained by projecting each canonical component onto the null space of the other components to eliminate cross-component effects. The exponents were averaged over time to give estimates of stability of the system along each component.

We also assessed the synchronicity of the two systems. Difference along 𝑖 th canonical components between two systems at time t, denoted by 𝑒^𝑖^, is expressed as

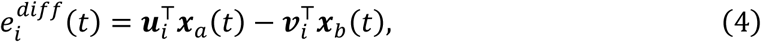

where 𝒖_𝑖_ and 𝒗_𝑖_ is canonical components of 𝒙_𝑎_ and 𝒙_𝑏_ respectively. The local Lyapunov exponent of 𝑒^𝑑𝑖𝑓𝑓^(𝑡) was estimated by perturbation experiments (described in “Directional perturbation experiment”). 𝑒^𝑑𝑖𝑓𝑓^(𝑡) was recorded for 1 sec after the perturbation and was averaged over at least 100 iterations. The mean data over time was fitted with exponential model 𝑦 = 𝑎 ∗ exp(𝜆𝑡) + 𝑐, with parameters 𝑎, 𝑐, and 𝜆. 𝑐 was introduced to allow for small baseline offset. The fitted coefficient 𝜆_𝑐𝑜ℎ,𝑖_ was used as an estimate of the local Lyapunov exponent of the 𝑒^𝑖^.

We thus obtain the DLIC index,

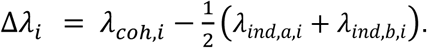

### Transient perturbation experiment

Transient perturbation experiment was done by applying random transient noise. The noise vector (512 dimensions) was sampled from the Gaussian distribution of mean 0 and standard deviation of 0.8. The noise vector was added to either RNN-A or -B randomly with probability of 0.005 per 10 ms.

### Directional perturbation experiment

Given a directional input v, it was normalized then multiplied by a random scalar chosen from Gaussian distribution of mean 0 and standard deviation 18. The standard deviation of 18 was chosen to match the expected norm of the noise vector used in the transient perturbation experiment. The noise vector was added to either RNN-A or -B randomly with probability of 0.005 per 10 ms.

### Computational resources

RNN training was executed on the TSUBAME 4.0 supercomputer (AMD EPYC 965 CPUs; NVIDIA H100 SXM5 94-GB GPUs). We ran the jobs on 72 “GPU-1” nodes in parallel, each node providing one H100 GPU and eight CPU cores 72 GPUs and 512 CPU cores in total).

### Statistics

Student t-test, Wilcoxon’s rank-sum test, chi-squared test, Spearman’s rank correlation coefficient was used for pairwise comparison. Analysis of variance (ANOVA) was used for multiple-comparison. Data was expressed as means ± S.E.M unless otherwise noted. The statistical significance of independent and consistent errors was assessed with a bootstrap procedure. First, for each trial we defined the offset as the difference between the decoded time and the true time. Offsets ≤ 1 s were considered correct; all others were labelled errors. An independent error occurred when one area (M2 or PPC) was correct, and the other was erroneous. A consistent error was counted when both areas were erroneous, but their offsets differed by ≤ 1 s. Let 𝑃 be the empirical probability that both areas were correct in a given session; then the total error probability is 1 − 𝑃, corresponding to (1 − 𝑃) × 𝑁 error points in a data set of 𝑁 trials. The joint offset distribution was binned into an 18 × 18 grid; the three-

by-three central region contained the correct trials, leaving 18 × 18 − 3 × 3 bins for errors. For each session we constructed a null distribution by randomly allocating the (1 − 𝑃) × 𝑁 error points among those error bins with equal probability, recomputing the rates of independent and consistent errors. This shuffling was repeated 1,000 times to generate the session-specific bootstrap distribution.

## Supplementary information

The supplementary information includes mathematical details on DLIC analysis and cPCA analysis.

## Code availability

The Python code for the RNN model will be released on GitHub upon the paper’s acceptance.

## Data availability

The experimental datasets will be made publicly available on Figshare once the manuscript is accepted for publication.

## Acknowledgements

We thank S. Kato and K. Kobayashi for AAV production, O. Ooishi for producing head-fixation apparatus, H. Kasai, K. Sakamoto, S.L. Smith, Y. Yu, T. Wakisaka for comments, and all members in Isomura lab for supporting this work. This work was supported by JP22wm0525007 (RH), JP25wm0625405 (RH) from AMED, JP22H02731 (RH), JP20K22678 (RH), JP21B304 (RH), JP21H05134 (RH), JP21H05135 (RH), JP21H0524 2(YI), and JP23H02589 (YI) from MEXT/JSPS, JPMJFR231X (RH), JPMJCR1751 (YI) from JST, Nakatani Foundation (RH), Shimadzu Foundation (RH), Takeda Science Foundation (RH), The Precise Measurement Technology Promotion Foundation (RH), Tateishi Science and Technology Foundation (RH), and Research Foundation for OptoScience and Technology (RH). This work utilized the TSUBAME4.0 supercomputer at the Institute of Science Tokyo for RNN training and analysis.

## Author contributions

H.I. and Riichiro H. conceived the project. H.I. F.I. conducted experiments and data analysis. H.I. designed and analysed the RNN model. Reiko H. conducted pilot experiments. Y.I. supervised the project. Riichiro H. H.I. wrote the paper with the comments of all authors.

---

**Extended Data Fig. 1.**
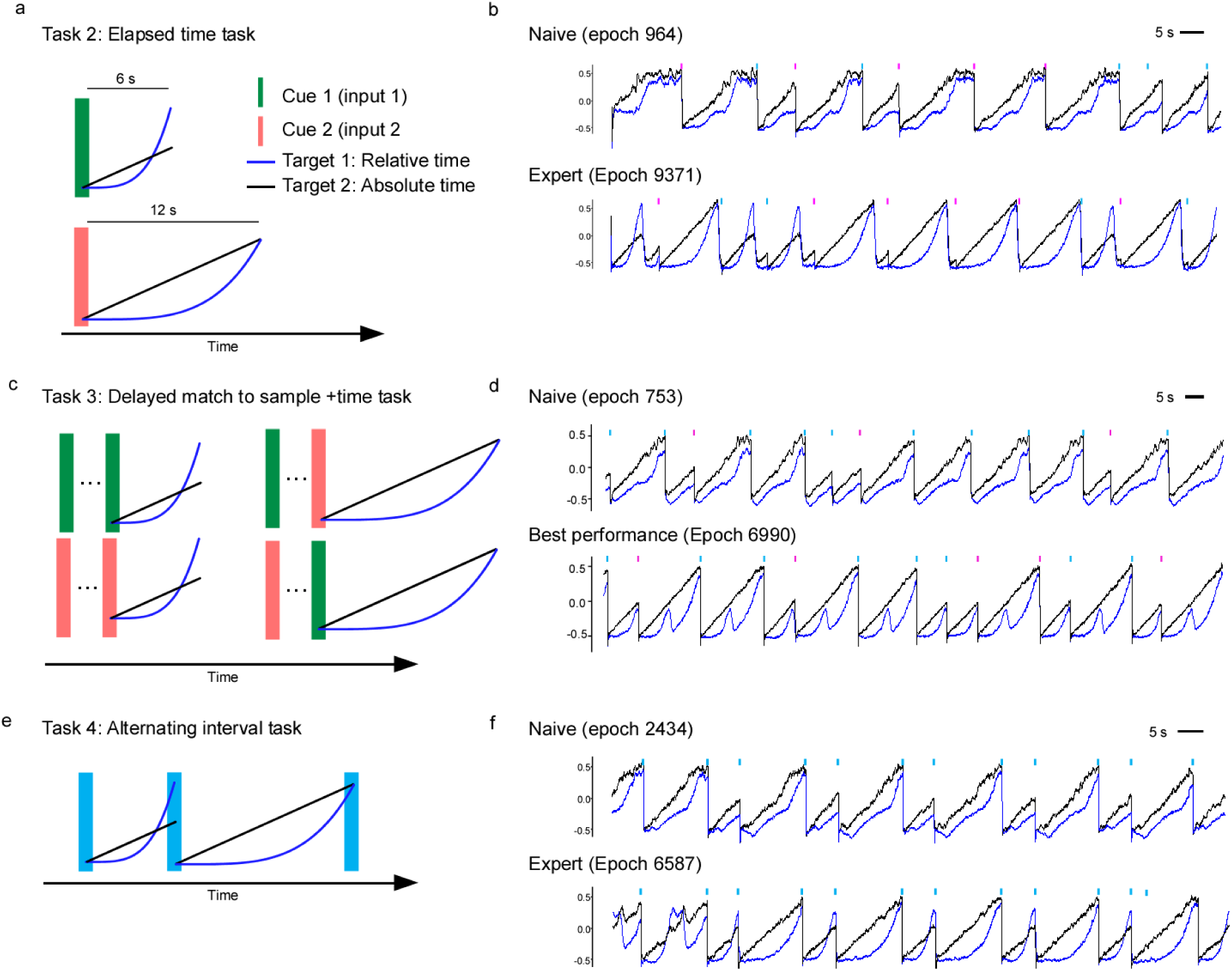
RNN training for three tasks. a. Two cues and their corresponding targets in the **elapsed time task**; Target 1 represents relative time, whereas Target 2 represents absolute time of the output durations. **b.** Example time-series outputs of the RNN in the elapsed time task: training in progress (top) vs. after training (bottom). **c.** Mapping between the order of the two preceding cues and the target in the **delayed match-to-sample + time task**. **d.** Example time-series outputs of the RNN in the delayed match-to-sample + time task: during training (top) vs. after training (bottom). Note that the agent still cannot generate appropriate outputs for relative time even when the network obtained the best score (epoch 9081). **e.** Relationship between cue order and target in the **alternating interval task**. Example time-series outputs of the RNN in the Alternating Interval task: during training (top) vs. after training (bottom).

**Extended Data Fig. 2.**
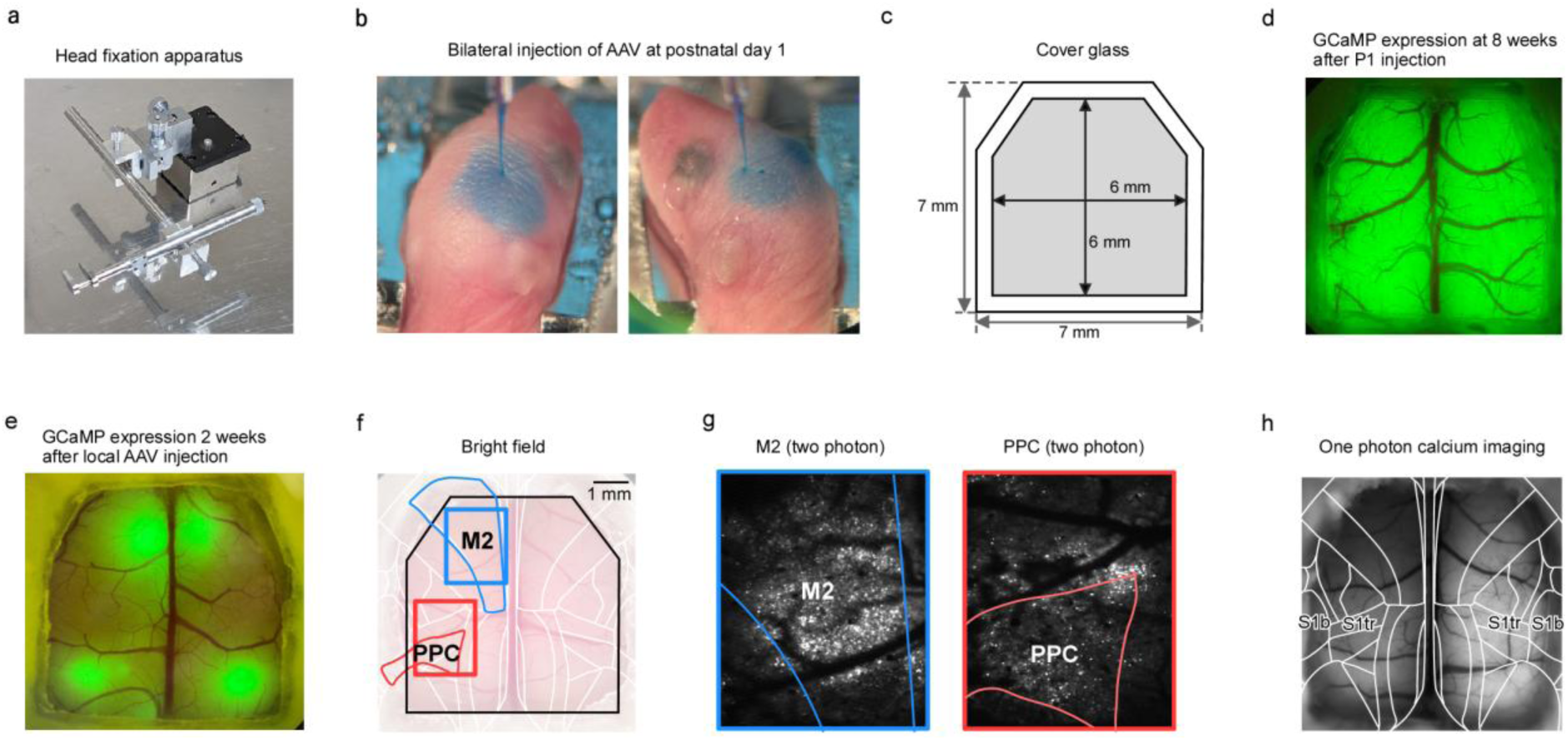
AAV injection and identification of cortical areas. a. Head-fixation apparatus for AAV injection in P1 mice. **b.** Example of bilateral AAV injection at P1. The spread of Fast Green is visible. **c.** Shape and size of the glass cranial window. **d.** Fluorescence image of a mouse with a cranial window eight weeks after P1 injection. **e.** Fluorescence image two weeks after four local AAV injections in an adult mouse. **f.** Example field of view for two-photon calcium imaging in a locally injected mouse. **g.** Two-photon calcium imaging image corresponding to panel **f**. Averaged image from one-photon calcium imaging showing the identified primary somatosensory area (trunk region) and S1 barrel cortex.

**Extended Data Fig. 3.**
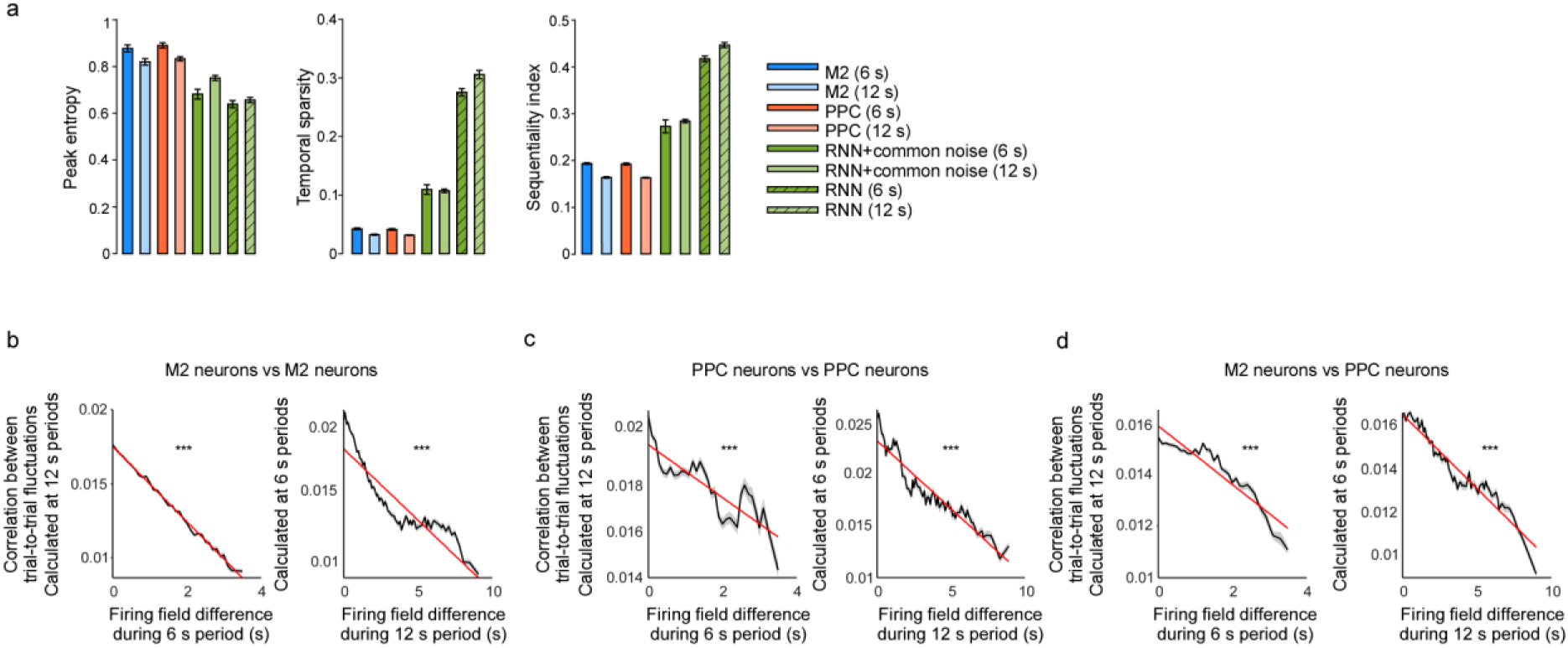
Comparison of M2 and PPC dynamics. a. The peak entropy, temporal sparsity, and sequentiality index were compared between M2, PPC, trained RNN, and trained noise-free RNN for both 6-s and 12-s periods. M2 versus PPC comparisons showed no significant differences for peak entropy (p = 0.094 at 6-s period, 0.16 at 12-s period), temporal sparsity (p = 0.094 at 6-s period, 0.44 at 12-s period), or the sequentiality index (p = 0.22 at 6-s period, 1.0 at 12-s period). **b.** For M2 neuron pairs, correlation between trial-to-trial fluctuations during the 12-s epoch were plotted against the differences in their firing-field times measured in the 6-s epoch (left). Conversely, correlations during the 6-s epoch were plotted against firing-field differences measured in the 12-s epoch (right). Linear fit (red line) and its slope are shown (*** p < 0.001, Pearson’s correlation test). **c.** The same analysis as **b** was conducted using PPC neuron pairs. The same analysis as **b** was conducted on pairs of M2 neuron and PPC neuron.

**Extended Data Fig. 4.**
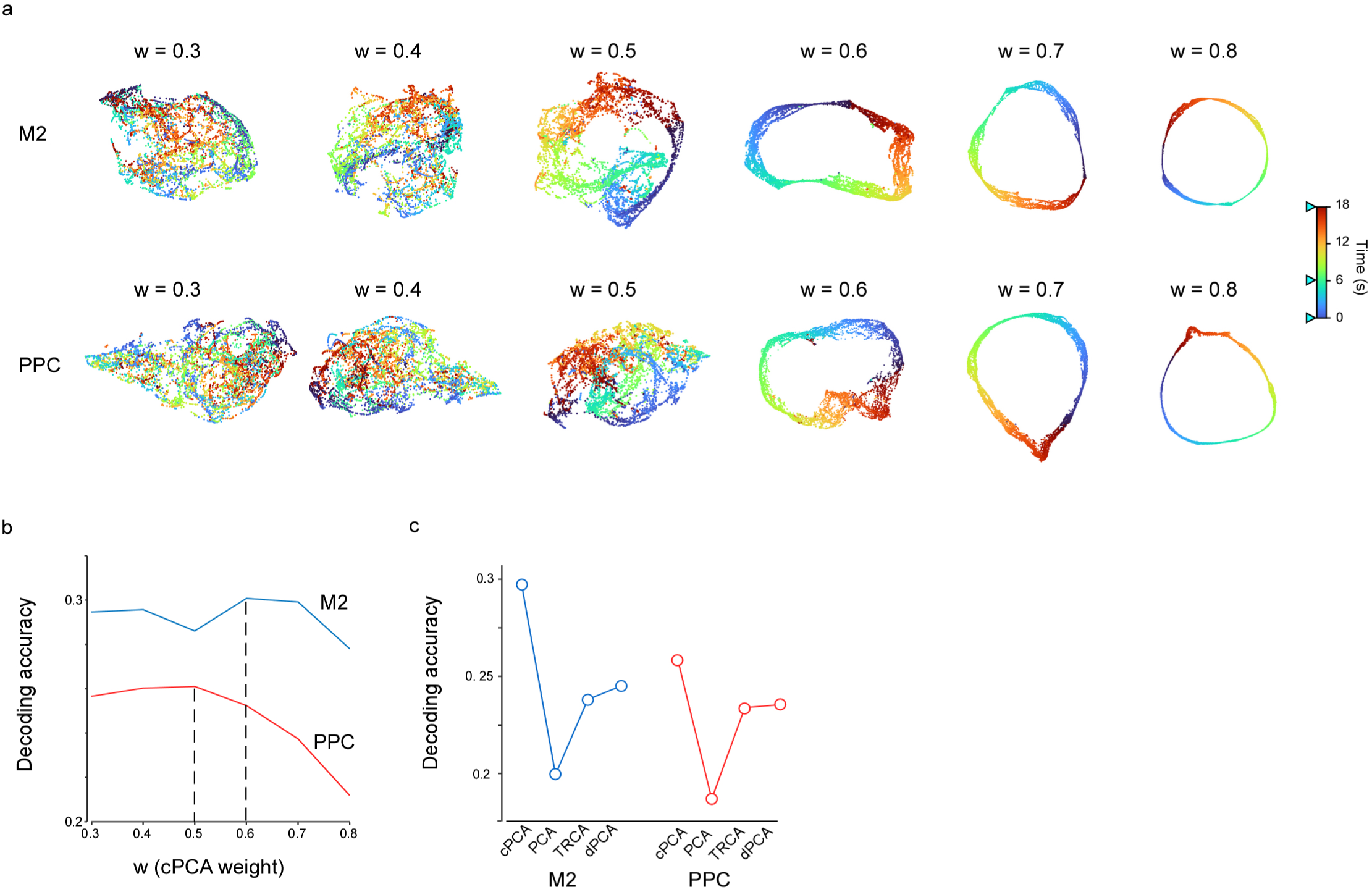
cPCA and decoding analysis. a. The population activity of M2 and PPC, averaged over every 5 trials, was dimensionally reduced using cPCA with the parameter w ranging from 0.3 to 0.8. UMAP was then applied to visualize the trajectory. **b.** Time-decoding performance of M2 and PPC plotted as a function of the cPCA weight parameter 𝑤. The decoding accuracy decreased at high 𝑤 values likely due to overfitting in both regions. Time-decoding accuracy with cPCA, PCA, TRCA, and dPCA.

**Extended Data Fig. 5.**
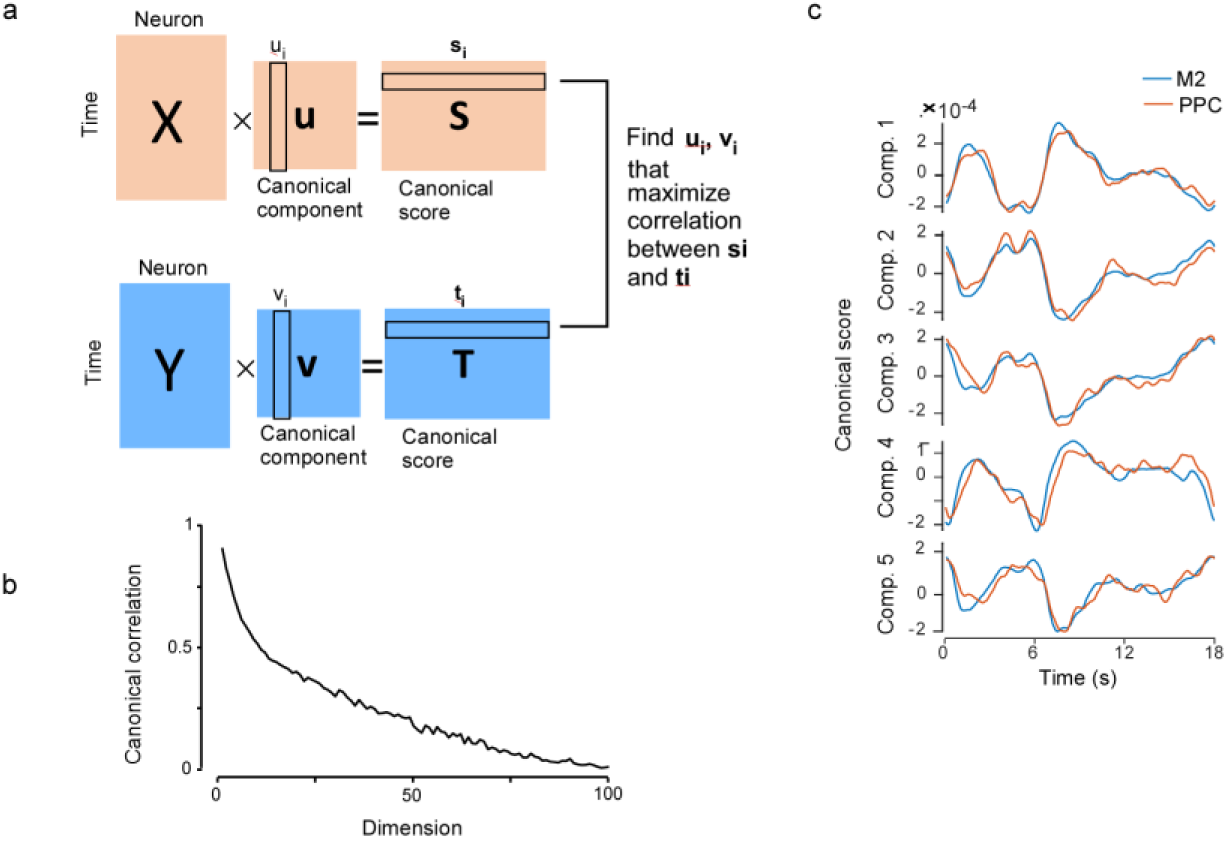
CCA analysis. a. Diagram illustrating CCA. **b.** Plot of canonical correlation as a function of dimension. Mean canonical scores over an 18-s period for M2 and PPC, shown for components 1–5.

**Extended data figure 6.**
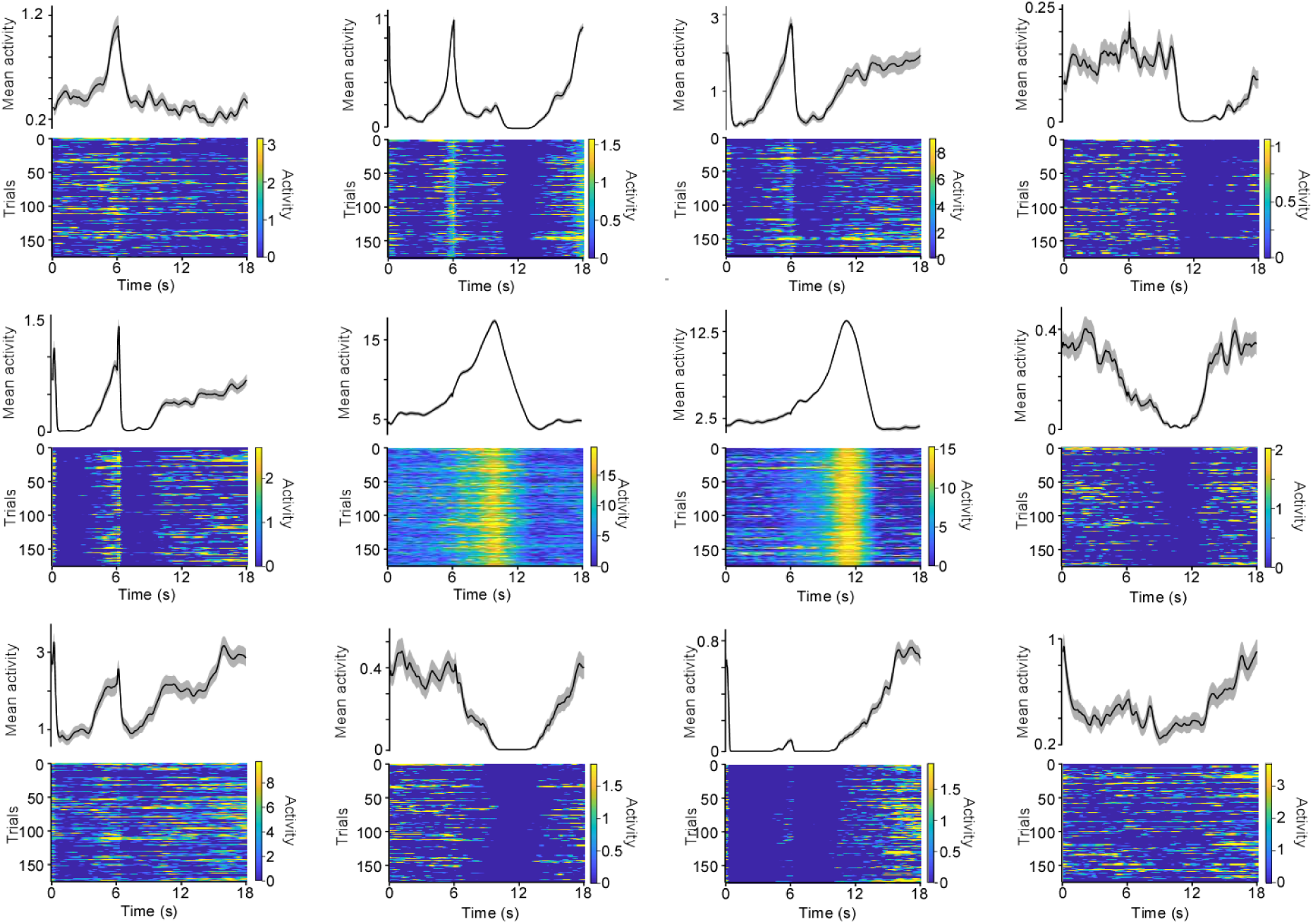
**Random examples of the unit activity of learned RNNs.** Mean activity and activity in each trial was shown for twelve units after learning which were randomly selected.

