## Supplementary information for "Independence and Coherence in Temporal Sequence Computation across the Fronto-Parietal Network"

This PDF file includes supplementary discussion for 1. DLIC analysis and 2. cPCA analysis.

### 1. DLIC analysis

#### Local Lyapunov exponent

**Local Lyapunov exponent.** We consider the bidirectionally–coupled dynamical system

$$\dot{\mathbf{x}}_a(t) = f\big(\mathbf{x}_a(t), \mathbf{x}_b(t)\big), \tag{1}$$

$$\dot{\mathbf{x}}_b(t) = g\big(\mathbf{x}_a(t), \mathbf{x}_b(t)\big), \tag{2}$$

where $\mathbf{x}_a, \mathbf{x}_b \in \mathbb{R}^{n\times 1}$ and a dot denotes differentiation with respect to time $t$.

Let $(\mathbf{u}_i, \mathbf{v}_i)$ be the $i$-th pair of canonical component vectors and define the scalar projection

$$c_a^{(i)}(t) = \mathbf{u}_i^\top \mathbf{x}_a(t) \tag{3}$$

$$c_b^{(i)}(t) = \mathbf{v}_i^\top \mathbf{x}_b(t) \tag{4}$$

**Perturbation dynamics.** Introduce an infinitesimal perturbation $(\delta\mathbf{x}_a, \delta\mathbf{x}_b)$ about the nominal trajectory. We first focus on system A. Linearising (3) gives

$$\delta c_a^{(i)}(t) = \mathbf{u}_i^\top \delta\mathbf{x}_a(t). \tag{5}$$

The full-state perturbation obeys the variational equation

$$\frac{\mathrm{d}}{\mathrm{d}t}\begin{pmatrix}\delta\mathbf{x}_a \\ \delta\mathbf{x}_b\end{pmatrix} = J(t)\begin{pmatrix}\delta\mathbf{x}_a \\ \delta\mathbf{x}_b\end{pmatrix}, \qquad J(t) = \begin{bmatrix}\dfrac{\partial f}{\partial \mathbf{x}_a} & \dfrac{\partial f}{\partial \mathbf{x}_b} \\ \dfrac{\partial g}{\partial \mathbf{x}_a} & \dfrac{\partial g}{\partial \mathbf{x}_b}\end{bmatrix}_{(\mathbf{x}_a(t),\mathbf{x}_b(t))}. \tag{6}$$

Differentiating (5) with respect to time and substituting the variational equation yields

$$\dot{\delta c}_a^{(i)}(t) = \mathbf{u}_i^\top \begin{pmatrix}\dfrac{\partial f}{\partial \mathbf{x}_a} & \dfrac{\partial f}{\partial \mathbf{x}_b}\end{pmatrix}\begin{pmatrix}\delta\mathbf{x}_a \\ \delta\mathbf{x}_b\end{pmatrix}. \tag{7}$$

If $\delta c_a^{(i)}(t) \approx Ae^{\lambda t}$ over a short window, the instantaneous (finite-time) local Lyapunov exponent along the $i$-th canonical direction is

$$\lambda_{a,i}(t) = \frac{\mathrm{d}}{\mathrm{d}t}\ln|\delta c_a^{(i)}(t)| = \frac{\dot{\delta c}_a^{(i)}(t)}{\delta c_a^{(i)}(t)}. \tag{8}$$

Negative values of $\lambda_i(t)$ indicate local contraction (stability), whereas positive values signal local expansion (instability) along the subspace spanned by $\mathbf{u}_i$ in system A.

**Lyapunov exponent along a canonical direction.** The mapping from the scalar perturbation $\delta c_a^{(i)}$ back to state–space increments $(\delta\mathbf{x}_a, \delta\mathbf{x}_b)$ is not unique. To isolate the stability of system A itself along $\mathbf{u}_i$, we constrain the perturbation to lie purely in that subspace:

$$\begin{pmatrix}\delta\mathbf{x}_a \\ \delta\mathbf{x}_b\end{pmatrix} = \begin{pmatrix}\mathbf{u}_i \\ 0\end{pmatrix}\delta z, \tag{9}$$

with scalar amplitude $\delta z$. Substituting (19) into (5)–(7) gives

$$\delta c_a^{(i)}(t) = (\mathbf{u}_i^\top \mathbf{u}_i)\,\delta z, \tag{10}$$

$$\delta \dot{c}_a^{(i)}(t) = \mathbf{u}_i^\top \frac{\partial f}{\partial \mathbf{x}_a}\,\mathbf{u}_i\,\delta z. \tag{11}$$

Dividing (21) by (20) yields the (independent) Local Lyapunov Exponent (LLE)

$$\lambda_{\text{ind},a,i}(t) = \frac{\mathbf{u}_i^\top \dfrac{\partial f}{\partial \mathbf{x}_a}\,\mathbf{u}_i}{\mathbf{u}_i^\top \mathbf{u}_i}. \tag{12}$$

Positive values indicate local divergence of the trajectory along $\mathbf{u}_i$, whereas negative values imply convergence. Similarly, the LLE of the system B along canonical component $\mathbf{v}_i$ is

$$\lambda_{\text{ind},b,i}(t) = \frac{\mathbf{v}_i^\top \dfrac{\partial g}{\partial \mathbf{x}_b}\,\mathbf{v}_i}{\mathbf{v}_i^\top \mathbf{v}_i}. \tag{13}$$

**Discrete-time implementation (Euler update).** In simulations, the continuous-time RNN is updated via a forward Euler step of size $\Delta t$:

$$\mathbf{x}_a(t+\Delta t) = \mathbf{x}_a(t) + \Delta t\, f\big(\mathbf{x}_a(t),\, \mathbf{x}_b(t)\big), \tag{14}$$

$$\mathbf{x}_b(t+\Delta t) = \mathbf{x}_b(t) + \Delta t\, g\big(\mathbf{x}_a(t),\, \mathbf{x}_b(t)\big). \tag{15}$$

Linearizing this map around the trajectory and projecting along $\mathbf{u}_i$ gives

$$\delta c_a^{(i)}(t+\Delta t) = \mathbf{u}_i^\top \left(I + \Delta t\,\frac{\partial f}{\partial \mathbf{x}_a}\right)\mathbf{u}_i\;\delta z, \tag{16}$$

so that the finite-time Lyapunov exponent reads

$$\lambda_{\text{ind},a,i}^{\Delta t}(t) = \frac{1}{\Delta t}\ln\left|\frac{\delta c_a^{(i)}(t+\Delta t)}{\delta c_a^{(i)}(t)}\right| = \frac{1}{\Delta t}\ln\left|\frac{\mathbf{u}_i^\top \left(I + \Delta t\,\dfrac{\partial f}{\partial \mathbf{x}_a}\right)\mathbf{u}_i}{\mathbf{u}_i^\top \mathbf{u}_i}\right|. \tag{17}$$

We evaluate (17) along a reference trajectory and average over a suitable time window to obtain the discrete-time LLE for the $i$-th canonical component.

### Coherence between the two subsystems

Canonical–correlation directions are expected to align the two states. To test synchrony we therefore examine the difference projection

$$e_i^{\text{diff}}(t) = \mathbf{u}_i^\top \mathbf{x}_a(t) - \mathbf{v}_i^\top \mathbf{x}_b(t). \tag{18}$$

The goal is to evaluate how fast $e_i^{\text{diff}}$ converges to 0 after perturbation by assuming an exponential relaxation $e_i^{\text{diff}} = Ae^{\lambda t}$. Because the nominal value of (18) is zero, its perturbation is simply $\delta e_i^{\text{diff}} = e_i^{\text{diff}}$. To obtain LLE, we constrain the perturbation to lie along axis that increases the canonical score difference.

$$\begin{pmatrix}\delta\mathbf{x}_a \\ \delta\mathbf{x}_b\end{pmatrix} = \begin{pmatrix}\mathbf{u}_i \\ -\mathbf{v}_i\end{pmatrix}\delta z, = \mathbf{w}_i\delta z \tag{19}$$

This yields

$$\delta e_i^{\text{diff}}(t) = (\mathbf{w}_i^\top \mathbf{w}_i)\,\delta z, \tag{20}$$

$$\delta \dot{e}_i^{\text{diff}}(t) = \mathbf{w}_i^\top J(t)\,\mathbf{w}_i\,\delta z. \tag{21}$$

Hence the coherence exponent is

$$\lambda_{\text{coh},i}(t) = \frac{\mathbf{w}_i^\top J(t)\,\mathbf{w}_i}{\mathbf{w}_i^\top \mathbf{w}_i}. \tag{22}$$

In the numerical evaluation, $e_i^{\text{diff}}$ was recorded following the perturbation along $\mathbf{w}_i$ and fitted with $Ae^{\lambda t} + c$, where $A, \lambda, c$ are parameters. $c$ was introduced to allow for a small baseline offset. Standard canonical component was used to ensure that the nominal value of $e_i^{\text{diff}}$ was centered at zero.

### DLIC index: relative synchrony

A large negative $\lambda_{\text{coh},i}$ can arise either because the two subsystems actually converge to each other or because both collapse rapidly along the same direction. To distinguish these possibilities we define the Dual LLE-based Independence vs Coherence (DLIC)

$$\Delta\lambda_i \;=\; \lambda_{\text{coh},i} - \frac{1}{2}\left(\lambda_{\text{ind},a,i} + \lambda_{\text{ind},b,i}\right). \tag{23}$$

- $\Delta\lambda_i < 0 \Longrightarrow$ the difference decays faster than the individual components, signalling genuine synchronisation.
- $\Delta\lambda_i > 0 \Longrightarrow$ residual desynchronisation along the $i$-th canonical axis.

Finally, using (12)–(22) and the block structure of $J(t)$ one finds the compact analytical form

$$\Delta\lambda_i = -\left(\mathbf{u}_i^\top \frac{\partial \dot{\mathbf{x}}_a}{\partial \mathbf{x}_b}\,\mathbf{v}_i + \mathbf{v}_i^\top \frac{\partial \dot{\mathbf{x}}_b}{\partial \mathbf{x}_a}\,\mathbf{u}_i\right). \tag{24}$$

with $\|\mathbf{u}_i\|^2 = \|\mathbf{v}_i\|^2 = 1$.

### Application of the analytical DLIC formula to the twin-RNN

**Twin-RNN dynamics.** The two RNNs obey

$$\tau\,\dot{\mathbf{x}}_a = -\mathbf{x}_a + \sigma\big(\mathbf{h}_a\big), \tag{25}$$

$$\tau\,\dot{\mathbf{x}}_b = -\mathbf{x}_b + \sigma\big(\mathbf{h}_b\big), \tag{26}$$

where

$$\mathbf{h}_a = W_a^{\text{RNN}}\mathbf{r}_a + S_b^{\text{RNN}}\mathbf{r}_b + W_a^{\text{IN}}\,u + \boldsymbol{\phi}_a, \tag{27}$$

$$\mathbf{h}_b = W_b^{\text{RNN}}\mathbf{r}_b + S_a^{\text{RNN}}\mathbf{r}_a + W_b^{\text{IN}}\,u + \boldsymbol{\phi}_b. \tag{28}$$

Here $\mathbf{x}_{a,b} \in \mathbb{R}^{N\times 1}$ are membrane potentials, $\mathbf{r}_{a,b} = \text{ReLU}(\mathbf{x}_{a,b})$ are firing rates, $u \in \mathbb{R}$ is a scalar input, $\boldsymbol{\phi}_{a,b}$ are zero-mean Gaussian noise vectors, $W^{\text{RNN}}$ are intra-network weights, and $S^{\text{RNN}}$ are inter-network weights. The nonlinear function is $\sigma(z) = \text{LeakyReLU}_\alpha(z)$ with slope $\alpha = 0.2$ for negative arguments.

**Jacobian.** Let $\Sigma_{a,b} = \text{diag}\big(\sigma'(\mathbf{h}_{a,b})\big)$ and $R_{a,b} = \text{diag}\big(\mathbf{1}_{\mathbf{x}_{a,b}>0}\big)$. The block Jacobian of the full system, based on (6), is

$$J(t) = \frac{1}{\tau}\begin{bmatrix} -I + \Sigma_a W_a^{\text{RNN}} R_a & \Sigma_a S_b^{\text{RNN}} R_b \\ \Sigma_b S_a^{\text{RNN}} R_a & -I + \Sigma_b W_b^{\text{RNN}} R_b \end{bmatrix}. \tag{29}$$

**Analytical DLIC.** Substituting (29) into the general formula (24) gives

$$\Delta\lambda_i(t) = -\frac{1}{\tau}\Big(\mathbf{u}_i^\top\,\Sigma_a S_b^{\mathrm{RNN}} R_b\,\mathbf{v}_i + \mathbf{v}_i^\top\,\Sigma_b S_a^{\mathrm{RNN}} R_a\,\mathbf{u}_i\Big). \tag{30}$$

**Time-averaged, activity-independent approximation.** $\Sigma_{a,b}$ and $R_{a,b}$ are diagonal matrices whose elements lie in $[\alpha, 1]$ and $\{0, 1\}$ respectively, and their entries fluctuate with network activity. A time-invariant approximation is obtained by replacing them with identity matrices,

$$\overline{\Delta\lambda_i} \;\approx\; -\frac{1}{\tau}\big(\mathbf{u}_i^\top S_b^{\mathrm{RNN}}\mathbf{v}_i + \mathbf{v}_i^\top S_a^{\mathrm{RNN}}\mathbf{u}_i\big). \tag{31}$$

This formula provides a meaningful interpretation.

- $\mathbf{u}_i^\top S_b^{\mathrm{RNN}}\mathbf{v}_i$ measures whether an $i$-th-mode displacement in network A, propagated through the A→B pathway, is cancelled (<0) or amplified (>0) by the matching displacement in network B.
- $\mathbf{v}_i^\top S_a^{\mathrm{RNN}}\mathbf{u}_i$ plays the symmetric role for perturbations initiated in network B and fed back to A.

Figure 6j shows that these two inner products are large and negative for the first two canonical components, consistent with the strongly negative DLIC obtained from the full numerical evaluation (30).
Each term in (31) indicates how the twin RNNs cancel perturbations along the $i$-th canonical component by its inter-network connections. The network has learned to cancel most effectively along the first two components, which is expected because they carry the largest shared noise. Altogether, our analysis explains how learning in the presence of common noise organizes the effect of perturbations along distinct axes, enabling the cortical computation to switch between independent and coherent modes.

### 2. cPCA analysis

Let the neural activity be arranged as a three–way tensor $X \in \mathbb{R}^{N_{\text{trials}}\times T\times C}$, where $N_{\text{trials}}$ is the number of repeated trials, $T$ the number of time samples per trial, and $C$ the number of recorded neurons. Contrastive principal-component analysis (cPCA) seeks a spatial axis $v \in \mathbb{R}^C$ that maximises structure that is consistent within a trial while suppressing structure that merely varies *across* trials.

**Objective.** For a weighting parameter $0 \le \omega \le 1$, cPCA solves

$$\max_{\|v\|^2=1}\Big[(1-\omega)\,v^\top C_{\mathrm{w}} v - \;\omega\,v^\top C_{\mathrm{b}} v\Big], \tag{32}$$

where

- $C_{\mathrm{w}}$ is the intra-trial covariance matrix, capturing variability that repeats within a trial;
- $C_{\mathrm{b}}$ is the inter-trial covariance matrix, capturing variability that differs between trials.

Separate the two variability sources by subtracting different means:

$$R_{\mathrm{w}}^{(r)}(t) = X_{r,t,:} - \frac{1}{T}\sum_{t'=1}^{T} X_{r,t',:},$$

$$R_{\mathrm{b}}^{(t)}(r) = X_{r,t,:} - \frac{1}{N_{\text{trials}}}\sum_{r'=1}^{N_{\text{trials}}} X_{r',t,:}.$$

Summing the outer products of these residuals yields

$$C_{\mathrm{w}} = \sum_{r=1}^{N_{\text{trials}}}\sum_{t=1}^{T} R_{\mathrm{w}}^{(r)}(t)\,R_{\mathrm{w}}^{(r)}(t)^\top,$$

$$C_{\mathrm{b}} = \sum_{t=1}^{T}\sum_{r=1}^{N_{\text{trials}}} R_{\mathrm{b}}^{(t)}(r)\,R_{\mathrm{b}}^{(t)}(r)^\top,$$

and each is normalized by its trace. Problem (32) is solved by the leading eigenvectors of the contrastive matrix

$$M \;=\; (1-\omega)\, C_{\mathrm{w}} - \omega\, C_{\mathrm{b}}.$$

These eigenvectors $v_1, v_2, \ldots$ form a low-dimensional orthonormal basis that emphasizes trial-consistent dynamics.

Projecting the data onto the $k$-th component gives

$$y_{r,t,k} \;=\; v_k^{\top} X_{r,t,:},$$

yielding trajectories that capture the dominant temporal structure within each trial while remaining reproducible across trials.
